# Characterizing endogenous delta oscillations in human MEG

**DOI:** 10.1101/2022.12.15.520554

**Authors:** Harish Gunasekaran, Leila Azizi, Virginie van Wassenhove, Sophie K. Herbst

## Abstract

Rhythmic activity in the delta frequency range (0.5 – 3 Hz) is a prominent feature of brain dynamics. Here, we examined whether spontaneous delta oscillations, as found in invasive recordings in awake animals, can be observed in non-invasive recordings pewwrformed in humans with magnetoencephalography (MEG). In humans, delta activity is commonly reported when processing rhythmic sensory inputs, with direct relationships to behaviour. However, rhythmic brain dynamics observed during rhythmic sensory stimulation cannot be interpreted as an *endogenous* oscillation. To test for endogenous delta oscillations we analysed human MEG data during rest. For comparison, we additionally analysed two conditions in which participants engaged in spontaneous finger tapping and silent counting, arguing that internally rhythmic behaviours could incite an otherwise silent neural oscillator. A novel set of analysis steps allowed us to show narrow spectral peaks in the delta frequency range in rest, and during overt and covert rhythmic activity. Additional analyses in the time domain revealed that only the resting state condition warranted an interpretation of these peaks as endogenously periodic neural dynamics. In sum, this work shows that using advanced signal processing techniques, it is possible to observe endogenous delta oscillations in non-invasive recordings of human brain dynamics.

## 1. Introduction

Delta-band activity is a prominent feature of neural dynamics traditionally observed during states of absence of consciousness, such as non-REM sleep in animals [1], [2] and humans [3]–[5]. Delta-band activity in the local field potential has also been reported during wakefulness in animals [6]–[8]. In humans, a number of brain regions including the frontal, temporal, and occipital areas as well as the hippocampus show delta-band activity [4], [9], [10], when recorded invasively with electrocorticography (ECoG) and intracranial electroencephalography (iEEG) in patients with treatment-resistant epilepsy.

Rhythmic brain dynamics provide a natural biological implementation of temporal structures, which have long been assigned a functional role in perception and cognition [11]– [13]. Delta-band activity (0.5 – 3 Hz) could be important in this respect, as its biophysical properties allow for the synchronization of larger networks of brain areas to a common temporal regime [14]–[16]. Seminal work in non-human primates has shown that delta oscillations emulate the temporal structure of sensory inputs by *entraining* to it [17], [18], [8], [19], [20]. Aligning the phase of slow oscillations to the temporal structure of external inputs tunes the excitability of the respective sensory areas to relevant inputs, locally modulating the spike rate of neurons [21], [22]. More globally, slow oscillations have also been shown to orchestrate an oscillatory hierarchy through phase-coupling [8], [23], [24].

The frequency range of the delta band also coheres well with natural rhythms that constrain auditory inputs like speech or music [25]–[27], and active sensing [28], [29]. Building on the work in non-human primates, an important body of research demonstrated that delta band activity measured in the human magneto- / and electro- encephalogram (M/EEG) can phase-lock to periodic inputs, surfacing as increased stimulus-brain coherence in auditory and motor areas [30]–[42], a mechanism termed neural entrainment.

Entrainment can implement a temporal structure in attention as postulated in the influential theory of dynamic attending [43], [44]. Crucially, entrainment in the narrow sense assumes an endogenous, physiological predisposition of the neural system to oscillate at particular frequency ranges, which can then emulate the exogenous input through phase shifts and phase alignment [45]–[48]. However, it is difficult to conclude on the existence of endogenous delta oscillations in the presence of periodic input signals, which passively drive brain activity leading to higher power in the same frequency range.

An active role of delta phase entrainment is suggested by its modulation through top-down influences such as the attended sensory modality [17], [31], [49], task demands [20], perceptual grouping [50], and hierarchical rhythmic structure of inputs [51]. Furthermore, entrainment can occur selectively to one frequency present in the input [40], and be sustained after the offset of the periodic stimulus [52], but see [53]15/05/2023 17:26:00 and still affect behaviour [52], [54], [55], for a review see [56]. Finally, previous studies have shown that entrainment scales with the strength of temporal predictions, surfacing as enhanced phase coherence in anticipation of temporally predictive input [32], [35], [42]. Also, tonic increases in delta amplitude, observed during the reading of numbers and mental calculation, and a working memory task [3], [57], [58], suggest that there are experimental conditions devoid from periodic stimulation that can enhance delta-band activity, favouring the existence of endogenous delta oscillations.

Still, there is a missing link between the invasively proven existence of spontaneous delta oscillations, and the delta-like signals revealed by non-invasive neuroimaging studies. The superposition of various signals in human M/EEG recordings makes it difficult to separate endogenously oscillatory from stimulus-evoked activity. This issue bears the risk of conflating pre-stimulus oscillatory signatures (i.e. delta phase) with post-stimulus evoked activity [59]– [61], thereby challenging the interpretation of the above-described stimulus-brain coherence as an endogenous oscillation. Few studies have tested for the presence of endogenous oscillatory activity in the delta band [62], [63], [35], see also [64] for a similar approach in the theta band, [36], and none have unequivocally shown the existence of an endogenous delta oscillation underlying the observed effects. Hence, to date we do not know whether delta phase locking observed in human M/EEG recordings truly reflects the entrainment of an endogenous oscillation [46], [48].

To start closing this gap, we set out to test the presence of endogenous oscillatory delta activity in human MEG signals, recorded at rest. It is however possible, that despite a pre-disposition to oscillate at a particular frequency, the brain does not spontaneously do so, and hence we would not see oscillatory activity in the resting state recordings. Periodic (or quasi-periodic) sensory input could incite an otherwise silent oscillator [36], [47], [65]. To avoid the above-described confound between endogenously periodic signals and the passive tracking of exogenous periodicities in sensory inputs, we additionally analysed MEG signals recorded while participants engaged in spontaneous overt or covert rhythmic behaviours. The frequency of such behaviours has been argued to reflect a stable internal prior [66]–[69], but see [70].

Currently, the state-of-the-art criterion for periodic brain activity, or oscillations, is to observe a peak in the power spectrum in a narrow frequency range [71], a definition most commonly used to study alpha oscillations. Peaks in the delta frequency range are not spontaneously visible from M/EEG spectra recorded during wake, due to the generally higher amplitudes at low frequencies, the 1/f property [72], [73]. Dedicated signal processing techniques can overcome this issue IRASA, [74].

From a set of data collected in the context of a different protocol [75], three conditions were selected, to reflect different degrees of participants’ engagement in spontaneous rhythmic behaviour: **(1) resting** (eyes open) – no rhythmic behaviour, **(2) spontaneous finger tapping** – overt engagement in rhythmic motor behaviour, and **(3) silent counting** – covert rhythmic behaviour. Using specifically adapted signal processing techniques, we were able to observe delta band peaks in the power spectrum in all three conditions, with meaningful topographies but heterogeneous peak frequencies. However, the interpretation of these peaks as strictly oscillatory remains somewhat questionable in the presence of evoked activity. In the resting condition, additional time-domain analyses [76] suggest the presence of endogenously periodic neural dynamics in the delta range.

## 2. Methods

### 2.1. Participants and data acquisition

MEG recordings of 22 right handed participants, recruited as part of a protocol assessing time perception [75], were used for this study. All participants had normal or corrected-to-normal vision without any known neurological or psychiatric disorders. The experimental protocol was approved by the local Ethics Committee on Human Research ’Comité de Protection des Personnes Sud-Est VI’ (protocol: CEA 100 049 / ID RCB: 2018-A02586-49), and all participants provided written informed consent in accordance with this protocol, and in in conformity with the Declaration of Helsinki (2018).

Two participants had noisy MEG data and two participants did not comply with the task, yielding a total of 18 participants for the final analysis (10 males; age = 26 years, SD = 5). The MEG data were collected using the whole-head Elekta Neuromag Vector View 306 MEG system (Neuromag Elekta LTD, Helsinki) in a magnetically shielded room at a sampling rate of 1000 Hz. The MEG system had 204 planar gradiometers and 102 magnetometers that measure the relative magnetic field strength (fT/cm) and absolute magnetic fields (fT), respectively. The direct current (DC) method was adopted during recording such that no high-pass filters were applied, to allow investigating low frequency components in the data. Horizontal and vertical electro-oculograms (EOG) and the electro-cardiogram (ECG) were recorded during the session. Participant’s head position was measured before each run by means of four head position indicator coils placed over the frontal and mastoid areas. For behavioural responses, participants pressed one button on a Fiber Optic Response Pad (Elekta), using the index finger of their right hand.

From the original data set, we selected three runs per participant (runs 1, 5, and 7 in 14 participants, runs 2, 4, and 6 in three participants, and runs 2, 7, and 6 in one participant due to a technical problem in run 4). The differences in run numbers were due to a change in the protocol after the initial participants.

### 2.2. Experimental conditions and tasks

The experiment (depicted in **Figure 1**) was presented through Psychtoolbox [77], [78] for Matlab R2016a. From the 12 conditions recorded in the original study (total duration: 34 minutes), we chose three experimental conditions for this analysis: **resting** (eyes open), **spontaneous finger tapping**, and **silent counting**. The three conditions were chosen to vary the level of rhythmic behaviour, in order to examine the presence of delta oscillatory activity in the brain recordings. During rest, the participants were supposedly not engaged in any rhythmic activity, and hence this condition was selected to examine the presence of spontaneous endogenous delta oscillations in the absence of rhythmic behaviour. In the spontaneous tapping condition, participants were actively engaged in *overtly* rhythmic motor behaviour, and we hypothesised that this condition was the most likely to result in a peak in the power spectrum at the individual tapping rate, possibly a signature of an endogenous neural oscillator. We also hypothesized that silent counting may engage participants’ auditory and articulatory systems in a *covertly* rhythmic manner, which should result in a peak in the power spectrum around the counting rate.

**Figure 1.**
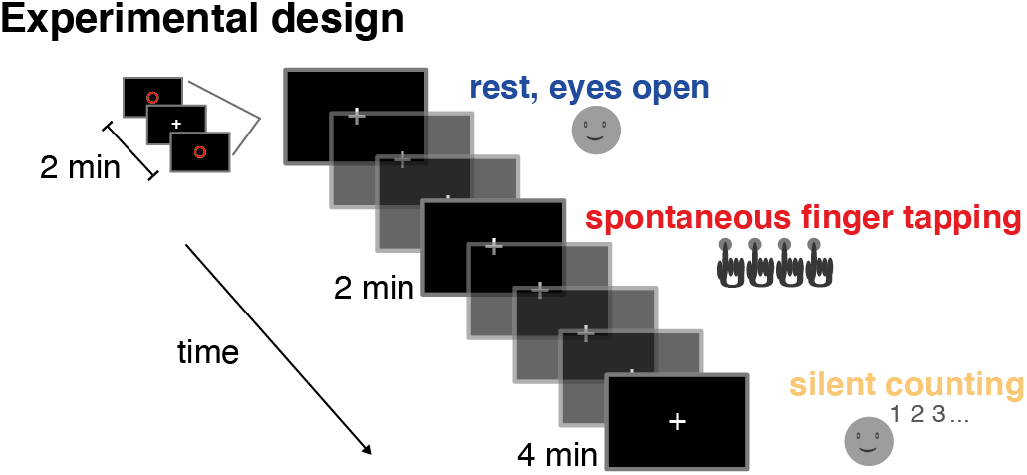
Experimental design. Experimental conditions selected from an existing dataset. We chose three conditions devoid of external stimulation, yet varying in the amount of internal rhythmic behaviour: Resting with eyes open (no rhythmic behaviour), spontaneous finger tapping (overt rhythmic behaviour), and silent counting (covert rhythmic behaviour).

In all runs, the beginning and end were marked by the French word "début" ("start") appearing on the screen for 1 s followed by a black screen lasting 4 s. The participant was instructed to start resting, tapping, or counting when a red circle briefly appeared (0.5 s) on the screen and until the same circle reappeared to mark the end of the run. Participants were instructed to steadily fixate the cross on the screen for the whole time and to sit as still as possible. During **resting** blocks, participants were informed that the aim was to record their brain activity at rest, and were only instructed to fixate the cross on the screen between the two occurrences of the red circle. **Spontaneous tapping** was executed on a tablet with the index finger of the right hand, at the individual’s own preferred pace. During **silent counting**, participants were asked to silently count without opening the mouth, and to report the final number after the run. Participants were not informed about the duration of each run.

During both tapping and counting, participants were additionally asked to estimate the duration of the run in seconds, resulting in a dual-task situation. The final duration had to be stated out loud to the experimenter by the interphone system. During rest, participants were not informed that they had to estimate duration. Four participants were asked for a retrospective duration estimate after the end of the run, but then the experimental protocol was changed. Due to this change in the experimental protocol, the runs were of slightly different duration, lasting 120 s for resting (300 s in the four participants for whom run 2 was used), 120 s for tapping (180 s if run 4 or 7 was used), and 240 s for counting (300 s if run 6 was used). As described below, we only used the first 120 s of MEG recordings from all runs.

### 2.3. Analyses of the behavioural data

#### 2.4.1. Behavioural tapping rate

Participants’ button presses were registered as time-stamped events with the MEG data, sampled at 1000 Hz. In accordance with the analyses of the MEG data (described below), the button-press time-series (coded as 0 when no button press occurred, and 1 when a press occurred) were subjected to a spectral analysis, by computing the Welch periodogram with a frequency resolution of 0.1 Hz, as implemented in MNE Python [79], [80]. Resulting power values were transformed to decibel (dB). A peak detection algorithm implemented in Scipy [81] was used to detect all peaks in the frequency range between 0.1 and 4 Hz. Then, the most prominent peak (according to scipy’s *peak_prominences* function) was retained as the behavioural tapping rate.

#### 2.4.2. Covert behavioural counting rate

Assuming that individuals silently counted, the covert individual counting frequencies were estimated by dividing the final number reported by the participant by the run’s duration (240 or 300 s). Since the counting was silent, we had no trace of the regularity of each individual’s counting. This measure was not available for one participant, who did not report a final count.

#### 2.4.3. Relative subjective duration estimates

To obtain the relative individual subjective duration estimates for the tapping and counting runs, we divided the duration in seconds reported by the participant by the duration of the corresponding run. This way, relative estimates below one reflect underestimation, and relative estimates above one reflect overestimation of the run’s duration.

### 2.4. MEG pre-processing

MEG data were processed using MNE python [79], [80]. The pre-processing consisted of the following steps: applying spatial signal source separation (SSS), low pass filtering, down-sampling, epoching, manual inspection and rejection of noisy epochs, and removal of ocular and cardiac artifacts using independent component analysis (ICA).

The raw data (DC, no high-pass filter was used) were corrected for environmental noise through spatial signal source separation, using the *Maxfilter* algorithm as implemented in MNE python, with the middle run (tapping) being used as a reference run to re-align the head coordinates. The algorithm also interpolates bad sensors, which are typically characterized by heavy distortions, flux jumps, baseline drifts, etc., and were identified through visual inspection for each individual.

Low-pass filtering was applied with a 100 Hz cut-off using a hamming windowed zero phase FIR filter. No high-pass filter was applied to avoid any signal distortion in the lower frequency ranges. The filtered data was down-sampled to 256 Hz. We then segmented the first 120 s of data from each run into 10 s long epochs with 8 s overlap, resulting in 168 epochs per participant. We subtracted the mean of each epoch for baseline correction, and applied linear detrending over the 10 s window. Noisy epochs were identified and rejected through visual inspection (21 epochs, in one participant only). For further analysis, we only used the 102 magnetometers.

Importantly, a well-known artifact of physiological origin apparent in human MEG, cardiac activity, lies in the same frequency range as the delta-band activity of interest. Therefore we applied an extended ICA procedure to ensure its complete removal. ICA was run in two steps: first, we ran one ICA on the epoched data of all conditions jointly, to identify and remove ocular artifacts, and second, we ran another ICA to identify cardiac artifacts. All ICAs were run on the epoched data, high-pass filtered at 1 Hz only for this purpose. For the detection of artefacts related to the electro-oculogram (EOG), we used an inbuilt routine in MNE python, which uses the signal from the external electrodes to estimate a participant’s typical EOG activity, and returns the ICA components that correlate with these typical events (threshold: 3.0, z-score).

In order to best identify and remove the cardiac artifacts, we then used the EOG-cleaned data, and ran a set of ICAs, iterating over the number of ICA components, a parameter that determines the partition of variance explained by each component. Inspection of the ICA results had revealed that when using the same partition of variance for each participant, residual cardiac artifacts were left in the data. More specifically, the number of components parameter determines the number of principal components (during a pre-whitening principal component analysis, PCA) that are passed to the ICA algorithm during fitting, and is given as a proportion of variance explained, from 0 – 1. We here parametrized the proportion between 0.849 – 0.999, in steps of 0.025. Components in the MEG data reflecting cardiac activity were estimated by computing the typical cardiac event from the external recordings of cardiac activity threshold 0.1, cross-trial phase statistics, [82]. For each decomposition, we then identified the components correlating best with the typical ECG event of each participant. To determine the best decomposition for each participant, we iterated across all versions of the data after running ICA and rejecting the cardiac components, and computed an *ECG score*: the power ratio at the individual ECG peak in the MEG data, before and after the removal of the ECG components. The individual ECG peak frequency was first determined based on the power spectrum computed directly on the ECG recording (Welch power spectral density). We then computed a spatial filter based on the peak’s topography in the MEG data (activity distribution across sensors at the peak frequency). The power ratio of this peak (dot product of all sensors multiplied with the spatial filter) was computed before and after component removal via ICA, and the ICA solution which yielded the greatest reduction in the cardiac peak topography was retained for the participant.

### 2.5. Spectral analyses

From the cleaned epochs we computed the power spectral density (PSD, see **Figure 2A** for an exemplary participant) using *Welch’s periodogram* with a frequency resolution of 0.1 Hz, resulting in power estimates at 1281 frequency values. The power spectrum for each magnetometer was obtained by averaging the power spectrums across all the epochs. Power spectra were cut at 45 Hz and transformed to decibel (dB).

**Figure 2.**
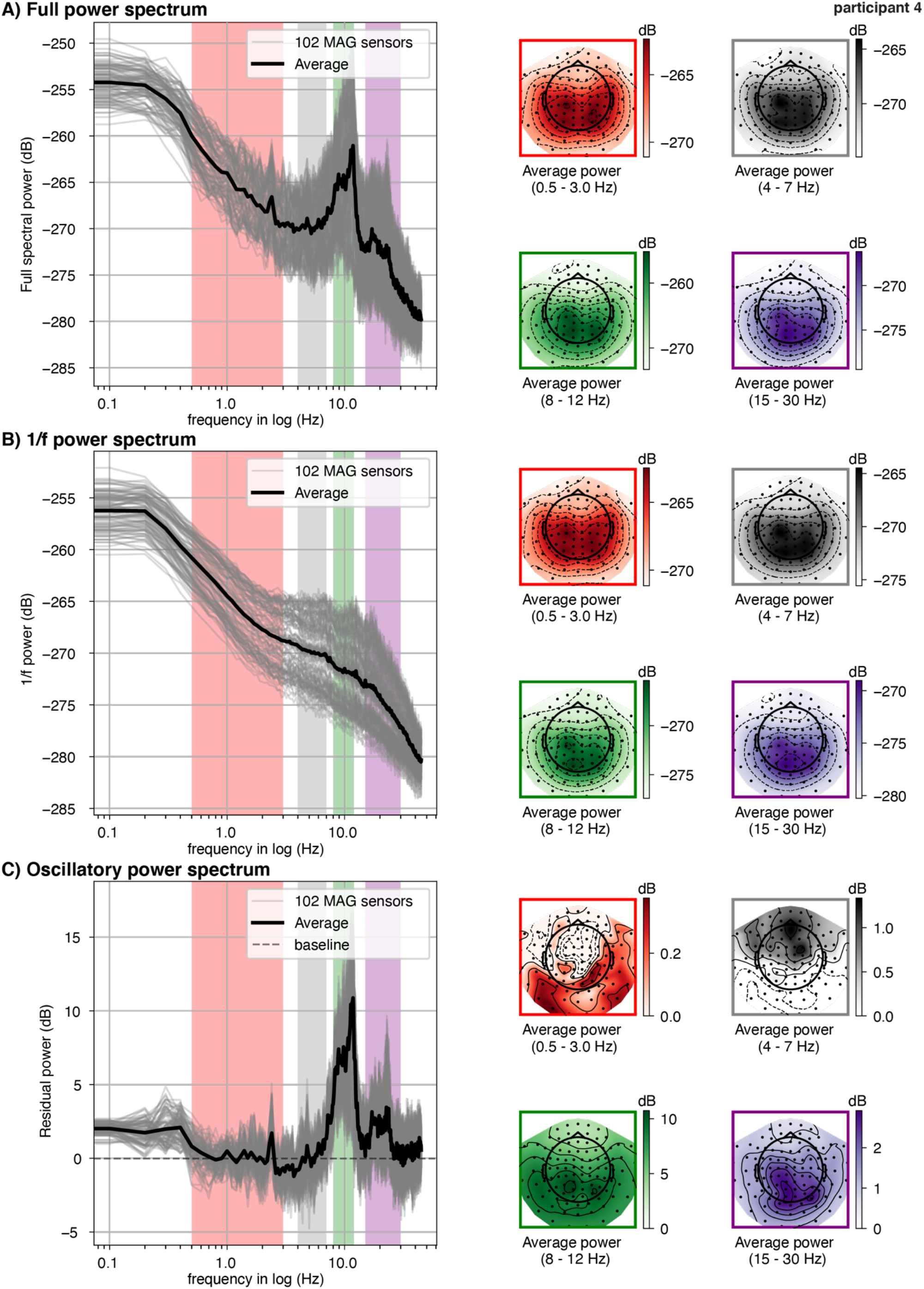
Exemplary power spectra from a single participant in the tapping condition. **A. Full power spectra** (0.1 – 45 Hz). Left: Welch’s power spectral density computed on 10 s long epochs, then averaged (grey lines depict the spectra of the 102 magnetometers, black line depicts average over sensors). The coloured shades depict the canonical frequency bands: delta (red, 0.5 – 3 Hz), theta (grey, 4 – 7 Hz), alpha (green, 8 – 12 Hz), beta (purple, 15 – 30 Hz). Right: Topographies averaged within in the canonical frequency bands. **B. 1/f power spectra.** Left: Power spectra computed on irregularly resampled data to obtain only the aperiodic components (IRASA method). Right: Topographies averaged in the canonical frequency bands. **C. Oscillatory power spectra**, obtained from subtracting the 1/f power spectra (depicted in B) from the full power spectra (depicted in A). Right: Topographies averaged in the canonical frequency bands.

A major concern when analysing delta band activity in M/EEG data, is that the peaks assumed to reflect narrow-band periodic activity lie in the high-power region of the 1/f aperiodic spectral component. To separate aperiodic spectral components from periodic ones, we applied the irregular resampling technique IRASA, [74]. The IRASA technique consists in down-sampling the epoched data in the time domain at pairwise non-integer values (here: 0.1 to 0.95 with a 0.05 increment), before computing the power spectrum. The 1/f power spectrum is obtained as the geometric mean of the power spectra at different resampling values (for an exemplary depiction see **Figure 2B)**.

To obtain the residual or oscillatory power spectra (**Figure 2C**), we subtracted for each participant and at each sensor the 1/f power spectrum from the full power spectrum. For further analysis, we divided the power spectrum into the canonical frequency bands, described in the literature [83], namely delta: 0.2 – 3.5 Hz, theta: 4 – 7 Hz, alpha: 8 – 12 Hz, beta: 15 – 30 Hz. The frequency range of delta activity was deliberately chosen slightly larger than the intended frequency band (0.5 – 3 Hz, with some variation in the literature) to allow the peak detection method described below to find peaks close to the desired cut-offs. **Figure 2** (right column) depicts the average topographies per canonical frequency band for an exemplary participant. Note the similarities in topographies between the full and 1/f spectrum especially in the delta band, which confirm the dominance of 1/f activity in the low frequency range. The theta, alpha, and beta bands were analysed similarly to the delta band, to validate the analysis pipeline on frequency bands for which peaks are more commonly reported in the literature.

### 2.6. Detection of spectral peaks

Per participant and condition, we identified spectral peaks in a given canonical frequency range. We assumed that if an oscillation is present, it should be reflected in higher power for a narrow range of frequencies at several sensors, while spurious peaks should vary in frequency across sensors. We thus extracted from each sensor the most prominent peak using peak finding algorithms implemented in Scipy [81], and applied k-means clustering as implemented in Scikit-learn see also [36], [84], to then identify peaks with coherent frequency across sensors (**see Figure 3**). While peaks with coherent frequency across sensors are more likely to reflect an underlying oscillation, this procedure does not provide a statistical test for oscillatory activity. The reason for not performing statistics at this point is that the peaks are defined per individual, and they are highly heterogenous, which does not allow for robust group statistics, as done for instance in the analyses SSVEP or ASSR (visual / auditory evoked steady state responses), when the expected peak frequency is known precisely.

**Figure 3.**
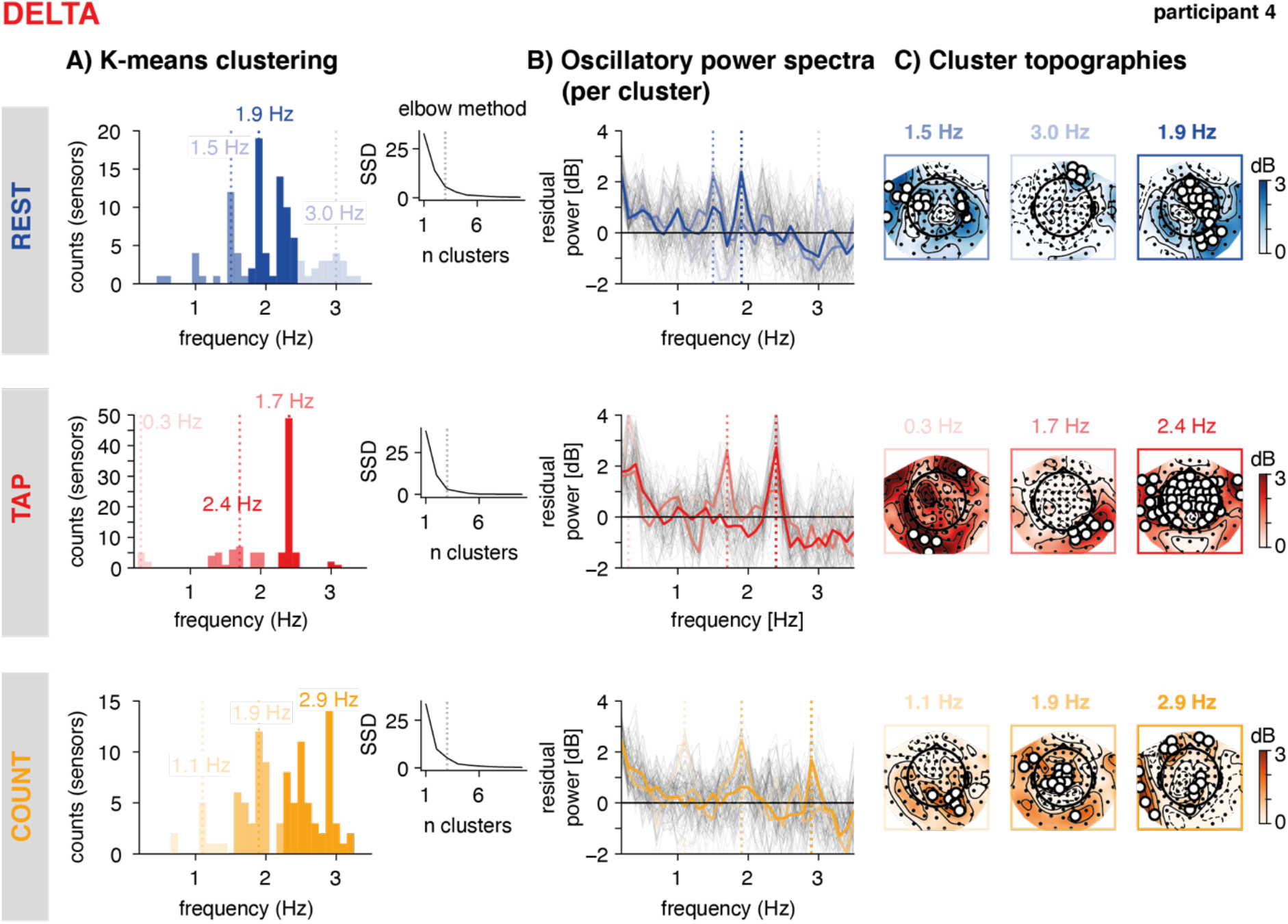
Peak finding method. Exemplary participant, delta-band. **A: K-means clustering.** Per condition (rest, tap, count, depicted as rows and in blue, red, and orange), spectral peaks identified per sensor were clustered using the k-means algorithm. Histograms depict the sensor counts per cluster, the brightness codes for cluster strength (brighter = higher), with the largest clusters (number of sensors included in cluster) in saturated colours. The vertical dashed lines indicate the peak frequency per cluster (defined by the mode). The number of clusters were determined from the data by identifying an ’elbow’ in the goodness of fit (SSD, the sum of squared distances from each point to its assigned cluster centre, see inset). **B: Oscillatory power spectra**, extracted from all sensors (grey lines), and the sensors in each cluster (coloured lines). The vertical dashed lines depict the peak frequency of each cluster. **C: Cluster topographies.** Power distribution across all sensors at the cluster’s peak frequency (± 0.1 Hz). The white circles depict the sensors selected as part of the cluster.

To obtain the peak frequency for each sensor, we applied a peak finding algorithm (scipy function *find_peaks*) which can result in several peaks, and in a second step computed peak prominences (scipy function *peak_prominence*) to select one peak per sensor as the one with maximal prominence. This resulted in 102 spectral peaks per canonical frequency band and condition, i.e. one per magnetometer. Once 102 peaks were identified for a given frequency band and condition, we applied k-means clustering of peak frequencies to identify the most consistent peak frequencies across sensors.

To determine the appropriate number of clusters, we used the so-called ’elbow method’ (depicted by the inset in **Figure 3A**), which consists in iterating over a range of possible n-clusters (here: 1 to 11), fitting the k-means algorithm for each n, and computing the inertia of the solution (automatically returned by Skicit-learn’s *KMeans* function). The inertia of the solution is a measure of goodness of fit of the solution, quantified as the sum of squared distances of samples to their closest cluster centre. The obtained vector of inertias over n-clusters was then examined for a flattening of the inertia values with higher n, i.e. an ’elbow’, using Skicit-learn’s *KneeLocator* function. Such a flattening reflects that the obtained solution does not improve strongly with higher n, and therefore the last n before the flattening is retained as the best and most parsimonious solution, and the k-means algorithm is fitted again with this n (average n across participants and conditions = 2.82, min = 2, max = 4, SD = 0.45). To characterize the clusters, we computed the **peak frequency** as the mode of the cluster (the frequency at which most of the contributing sensors showed a peak), the **peak power** as the average power over the contributing sensors at the peak frequency, as well as the **strength of the peak** as the number of sensors contributing to the cluster. The same method was applied to the theta, alpha, and beta bands (see **Figures S1, S2, S3**).

### 2.7. Peak sorting

The k-means clustering procedure produced several clusters per canonical frequency band, identified by their peak frequency, and ordered by cluster size (i.e., the number of sensors showing a peak at or close to this frequency). Depending on the frequency band at hand, we made different assumptions to group clusters across participants for further analysis.

#### 2.8.1. Delta band

We reasoned that the individual behavioural tapping frequency (see **Figure 4**) might result from an endogenously present oscillation at that frequency, or else, that spectral (MEG) peaks in the tapping condition reflect the periodically reoccurring tapping evoked response. Spectral peaks in the vicinity of the individual tapping rate were found for all participants, but varied in strength (hence cluster order), which prevented us from simply retaining the first cluster in the delta band. We thus identified for each participant one peak in the delta range (from the two to four peaks found by the k-means clustering) that was **closest to the individual tapping rate**. We further hypothesized that the individual tapping rate could reflect the stable frequency of an internal oscillator, which should then also be apparent in the resting and counting conditions in the absence of overt motor tapping. Thus we also identified peaks close to the individual *tapping* rate from the resting and counting runs. A second, **higher delta peak** was identified for all conditions, lying above the individual tapping rate. Here, we only selected peaks with higher frequencies, because for most participants the behavioural tapping rate was in the lower range of the delta band. A higher peak was not found in one participant in the tapping condition, and three participants in the resting condition, for which we then retreated to selecting the strongest cluster after removal of the tapping peak. Third, we identified the **peak closest to the individual counting rate** from all three runs. In sum, for delta, three types of peaks were obtained: the one **closest to tapping**, **high delta,** and **closest to counting** (see **Figure 5, Table 1**).

**Figure 4.**
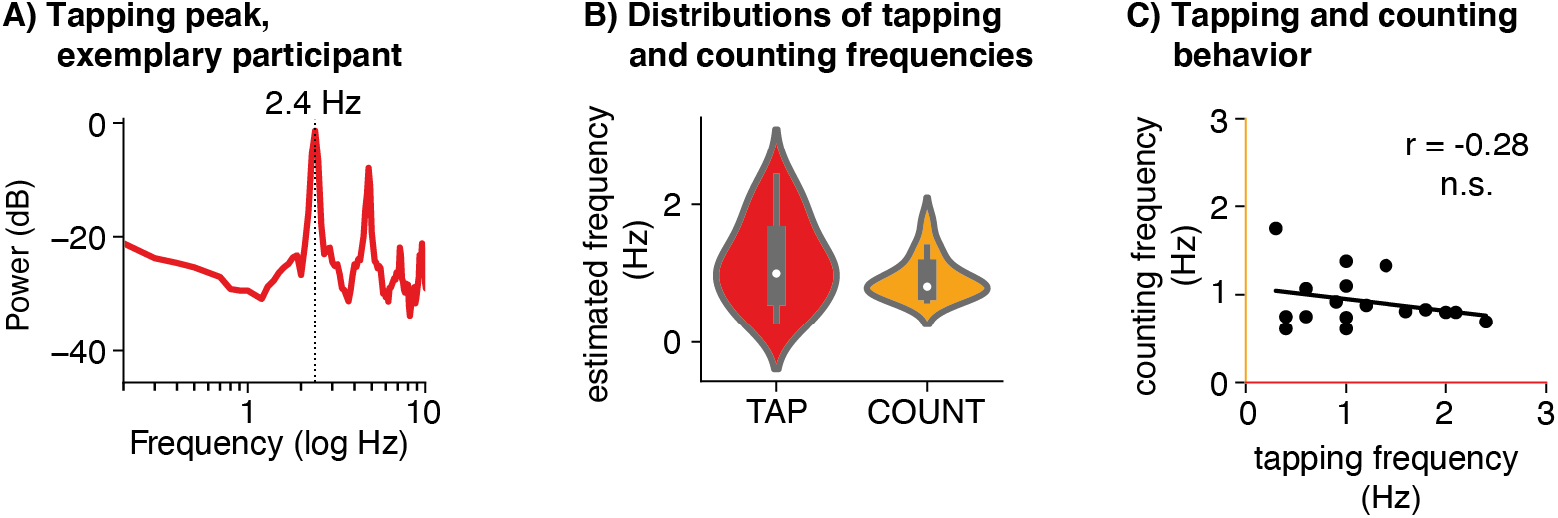
Behavioural data (N = 18). **A. Power spectral density of tapping behaviour,** depicted for an exemplary participant (same participant as in Figures 2 and 3). The vertical dashed line depicts the most prominent peak, indicating a tapping rate of 2.4 Hz. The peaks at higher frequencies reflect harmonics of the lowest peak. **B. Distributions of tapping and counting frequencies across participants.** Violin plots depict the distribution (kernel density estimation) of tapping (red) and counting (orange) frequencies across participants. White dots: median. Thick grey bar: interquartile range. **C. Tapping versus counting frequencies.** No significant correlation was found between individuals’ overt and covert behavioural rates.

**Figure 5.**
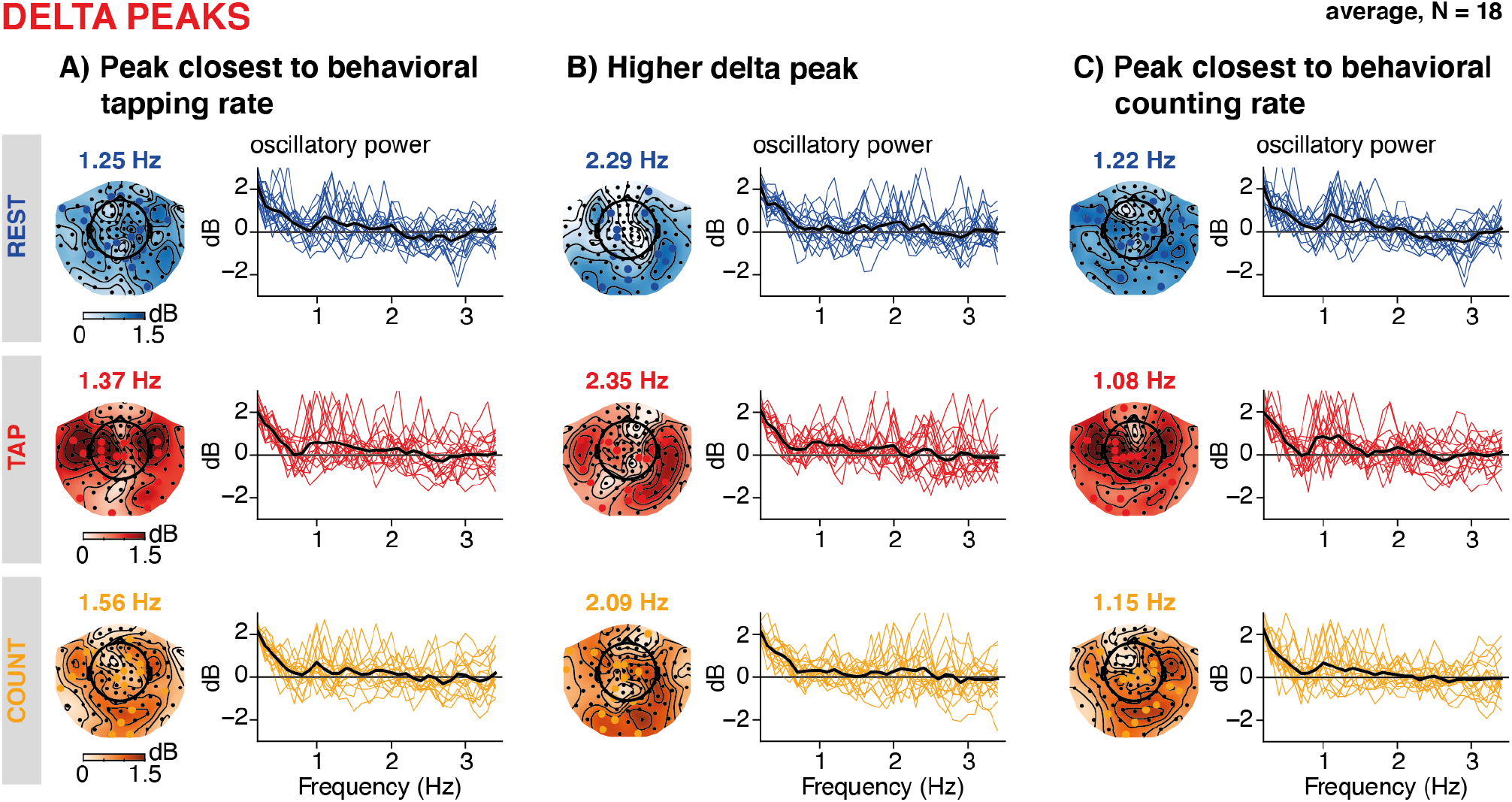
Result of the peak sorting procedure, delta band. A: Peak closest to behavioural tapping rates. The topographies show the power distribution at the individually identified peak frequencies, averaged across participants for the resting (top row, blue), tapping (middle row, red), and counting (bottom row, orange) conditions. The sensors marked in blue, red, or orange on the topographies depict individual peak sensors. The line plots show single participants’ oscillatory power spectral density averaged across all sensors in the cluster (coloured lines) and averaged across participants (black line). **B: Higher delta peaks.** A second cluster was identified as having a frequency above the individual tapping peak. **C: Peaks closest to the behavioural counting rates.** Independently from the peaks displayed in A and B, we also grouped peaks closest to the individual counting rates.

**Table 1:** Results of the peak clustering and sorting procedure. Different peaks were identified per canonical frequency band (delta, theta, alpha, beta, left column) and condition. The table indicates the average peak frequency across participants, as well as its standard deviation (SD), average power, its standard deviation, as well as number of sensors per cluster and its SD.

| freq. band | peak type | condition | freq. (mean, Hz) | freq. (SD, Hz) | power (mean, dB) | power (SD, dB) | number of sensors (mean) | number of sensors (SD) |
| --- | --- | --- | --- | --- | --- | --- | --- | --- |
| **DELTA** | tapping rate | rest | 1.25 | 0.43 | 3 | 0.64 | 11.06 | 5.07 |
|  |  | tap | 1.37 | 0.5 | 3.69 | 1 | 18.11 | 12.94 |
|  |  | count | 1.56 | 0.56 | 2.75 | 0.58 | 13.61 | 5.21 |
|  | high delta | rest | 2.29 | 0.5 | 2.97 | 1.04 | 13.67 | 5.04 |
|  |  | tap | 2.35 | 0.53 | 3.05 | 1.06 | 12.72 | 6.02 |
|  |  | count | 2.09 | 0.57 | 3.51 | 1.29 | 14.94 | 9.85 |
|  | counting rate | rest | 1.22 | 0.32 | 3.22 | 0.73 | 11.47 | 4.13 |
|  |  | tap | 1.08 | 0.32 | 3.33 | 0.96 | 11.41 | 7.03 |
|  |  | count | 1.15 | 0.33 | 2.97 | 0.65 | 11.76 | 4.58 |
| **THETA** | first | rest | 5.29 | 0.59 | 3.55 | 0.82 | 14.39 | 5.19 |
|  |  | tap | 5.38 | 0.54 | 3.89 | 1.94 | 17.61 | 9.6 |
|  |  | count | 5.31 | 0.35 | 3.55 | 0.81 | 15.78 | 5.73 |
| **ALPHA** | first | rest | 10.33 | 0.8 | 12.46 | 4.53 | 24.89 | 13.36 |
|  |  | tap | 10.48 | 0.62 | 11.75 | 4.28 | 22.17 | 10.47 |
|  |  | count | 10.18 | 0.84 | 12.32 | 4.75 | 28.44 | 14.6 |
| **BETA** | low | rest | 19.53 | 2.01 | 5.74 | 2 | 11.56 | 9.96 |
|  |  | tap | 18.96 | 2.13 | 6.23 | 2.27 | 9.61 | 4.5 |
|  |  | count | 19.16 | 2.49 | 5.43 | 1.82 | 8.89 | 4.79 |
|  | high | rest | 23.75 | 2.03 | 5.37 | 2.01 | 9.06 | 4.7 |
|  |  | tap | 23.57 | 1.92 | 5.12 | 2.38 | 10.17 | 4.51 |
|  |  | count | 23.24 | 2.08 | 5.44 | 2.5 | 9.78 | 4.06 |

#### 2.8.2. Theta and alpha bands

For both the theta and the alpha bands, we retained the **first** cluster (largest number of sensors) as the peak (see **Figure S4, S5, Table 1**), as these frequency-bands were not of primary interest to the study.

#### 2.8.3. Beta band

In the beta band, visual inspection showed that most participants’ first two clusters captured two distinct frequency ranges, namely a lower beta, and a higher beta cluster. As the strength varied between those two clusters across participants, we grouped the first two clusters for each participant into **low beta** and **high beta** peaks (**Figure S4, S5 Table 1**).

### 2.8. Statistical comparisons between conditions

To assess differences in delta-band activity between experimental conditions, we tested for condition differences in peak frequency, power, and the number of sensors contributing to a peak (**Figure 6, Table 2**). Paired two-sided t-tests were performed for all frequency bands and peak types, with a critical threshold of p < 0.01 to account for multiple testing.

**Figure 6.**
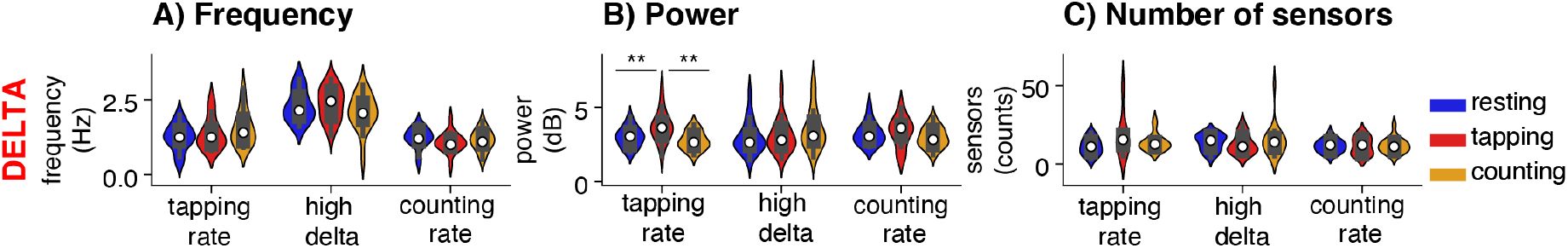
Peak frequency and power in the delta band across conditions. **A: Peak frequency.** Violin plots depict the distribution (kernel density estimation) of peak frequencies across participants. White dots: median. Thick grey bar: interquartile range. Frequency distributions are depicted for the peak selected as being closest to the individual behavioural tapping rate, the higher delta peak, and the peak at the counting rate. **B. Peak power.** As in A, but the y-axis reflects power. **C. Number of sensors in cluster.** As in A, but the y-axis reflects the number of sensors showing a peak at or close to this frequency. Significance values are indicated as follows: ** p < 0.01.

**Table 2.** Statistical tests of peak frequency, peak power, and number of sensors between conditions per canonical frequency band (DF = degrees of freedom).

| freq. band | peak type | cond 1 | cond 2 | peak frequency |  | peak power |  | number of sensors |  | DF |
| --- | --- | --- | --- | --- | --- | --- | --- | --- | --- | --- |
|  |  |  |  | T | P | T | P | T | P |  |
| **DELTA** | tapping rate | rest | tap | -1.11 | 0.284 | **-3** | **0.008** | -2.52 | 0.022 | 17 |
|  |  | rest | count | -2.49 | 0.023 | 1.38 | 0.186 | -1.43 | 0.171 | 17 |
|  |  | tap | count | -2.05 | 0.056 | **3.49** | **0.003** | 1.21 | 0.242 | 17 |
|  | higher | rest | tap | -0.39 | 0.702 | -0.26 | 0.8 | 0.68 | 0.503 | 17 |
|  |  | rest | count | 1.01 | 0.328 | -1.75 | 0.098 | -0.49 | 0.629 | 17 |
|  |  | tap | count | 1.32 | 0.205 | -1.25 | 0.228 | -0.76 | 0.459 | 17 |
|  | counting rate | rest | tap | 1.29 | 0.216 | -0.35 | 0.732 | 0.04 | 0.970 | 16 |
|  |  | rest | count | 0.75 | 0.465 | 1.56 | 0.139 | -0.21 | 0.835 | 16 |
|  |  | tap | count | -0.78 | 0.446 | 1.2 | 0.249 | -0.27 | 0.793 | 16 |
| **THETA** | first | rest | tap | -0.53 | 0.606 | -0.75 | 0.466 | -1.32 | 0.203 | 17 |
|  |  | rest | count | -0.14 | 0.891 | 0.02 | 0.987 | -0.74 | 0.472 | 17 |
|  |  | tap | count | 0.47 | 0.645 | 0.91 | 0.375 | 0.68 | 0.503 | 17 |
| **ALPHA** | first | rest | tap | -1.07 | 0.301 | 0.8 | 0.436 | 0.91 | 0.375 | 17 |
|  |  | rest | count | 1.43 | 0.17 | 0.18 | 0.857 | -0.85 | 0.41 | 17 |
|  |  | tap | count | 1.91 | 0.074 | -0.7 | 0.492 | -2.37 | 0.03 | 17 |
| **BETA** | low | rest | tap | 1.06 | 0.303 | -0.97 | 0.347 | 0.71 | 0.489 | 17 |
|  |  | rest | count | 0.57 | 0.574 | 0.69 | 0.502 | 1.05 | 0.31 | 17 |
|  |  | tap | count | -0.41 | 0.69 | 2.34 | 0.032 | 0.48 | 0.639 | 17 |
|  | high | rest | tap | 0.39 | 0.699 | 0.56 | 0.582 | -0.83 | 0.42 | 17 |
|  |  | rest | count | 0.89 | 0.384 | -0.15 | 0.883 | -0.49 | 0.63 | 17 |
|  |  | tap | count | 0.63 | 0.539 | -0.92 | 0.372 | 0.38 | 0.711 | 17 |

### 2.9. Correlation analyses

We were interested in whether delta-band peak frequencies correlate between conditions, or with the behavioural signatures, notably tapping rate. Simply correlating the peak frequencies would be trivial, as we had selected the delta peaks by their closeness to the behavioural tapping rate, hence introducing a strong correlation between behavioural rates and the peak frequencies. Instead of correlating the frequency values, we thus computed residuals from the linear fit between the peak frequency and the behavioural tapping rate, and respectively also the counting rate (in Hz). To compare the residuals between conditions, we used paired two-sided t-tests, as well as Pearson correlation coefficients (significance tests for correlations were performed with respect to Student’s t-distribution, with N−2 degrees of freedom) (**Figure 7**).

**Figure 7.**
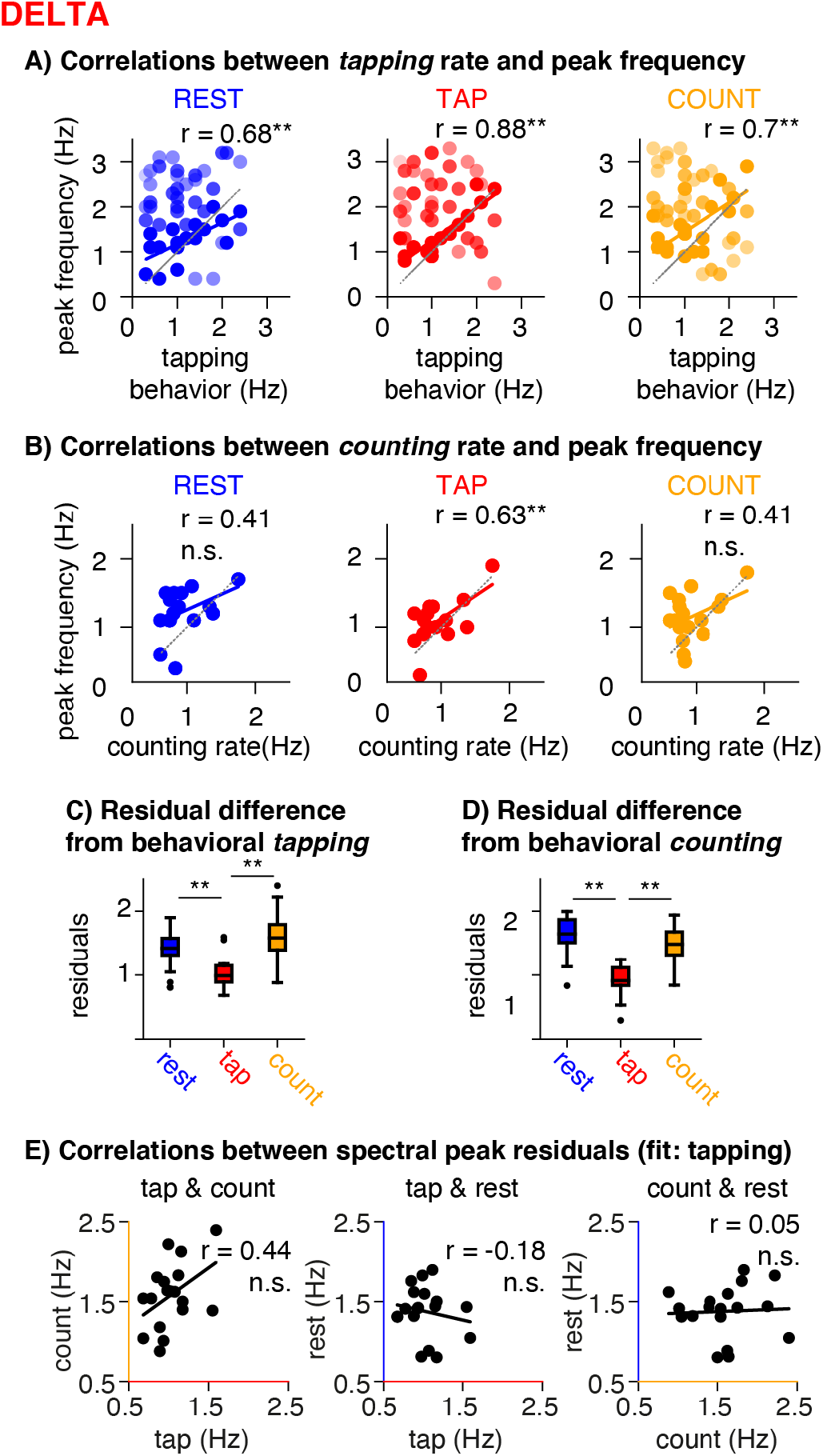
Correlation analyses (delta band). A. Correlations between the individual tapping rate and the frequencies of the spectral peaks identified in the delta band, per condition. The coloured line indicates the estimated correlation and the grey line the unity-line. Each dot is one cluster per participant with brighter colours indicating stronger clusters. The number of clusters identified varied over participants, hence the number of dots vary between panels. Significance values are indicated as follows: * p < 0.05; ** p < 0.01; n.s. not significant. B. Correlations between the individual counting rate and the frequencies of the spectral peaks. Same as in A, but with respect to the behavioural counting rate. C. Residual differences between the frequency of the spectral peak identified as nearest to the individual tapping rate, and the behavioural tapping rate per condition. Residuals were significantly smaller in the tapping condition compared to resting and counting. D. Residual differences between the frequency of the spectral peak identified as nearest to the individual counting rate, and the behavioural counting rate per condition. Residuals were smallest in tapping. E. Pearson correlations between the spectral peak residuals, that is the frequency of the spectral peak identified as nearest to the individual tapping rate after accounting for the correlations shown in A, for pair-wise combinations of conditions. No significant correlations were found between conditions.

### 2.10. Sensitivity analysis

Likely, the relatively small sample size (N = 18) limited our ability to detect significant differences and correlations [85]. A sensitivity analysis using G-Power [86] indeed indicated that only correlations above 0.61 (0.55 for one-sided tests), and mean differences with an effect size of 0.89 (Cohen’s d) could be detected with 80% power.

### 2.11. Tapping evoked responses

A spectral peak in the tapping condition would likely already result from the periodic occurrence of the sensorimotor evoked responses during tapping itself [87], [88]. To analyse the evoked response (see **Figure 8**) we epoched the data from the tapping conditions, time-locked to the registered button presses (−2 to 4 s), baseline corrected (−2 to 0 s) and detrended the epochs linearly over the full window. We then applied the ICAs computed for all runs to these epochs and rejected the previously identified ocular and cardiac components. To compute the peak topography, we averaged the epochs for each participant and applied Scipy’s peak detection algorithm in the window of 0 s to 0.25 s post-tap to extract the largest peak and compute its peak topography. We then correlated this topography (one amplitude value per sensor) with the peak topography at the delta peaks for each participant to test for spatial overlap.

**Figure 8.**
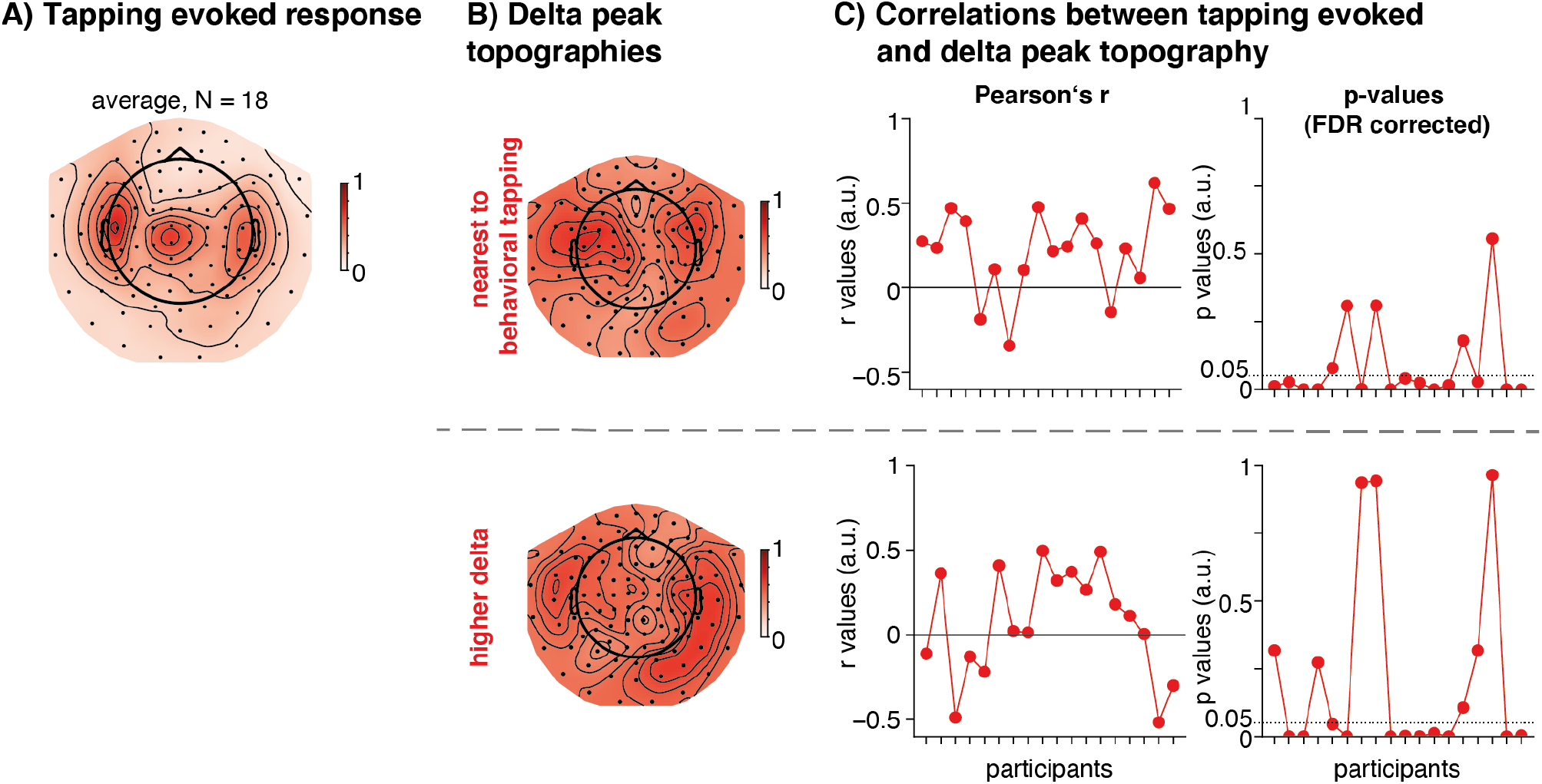
Topographies of the average tapping evoked response, and the average delta peak. **A. Average tapping evoked response** (peak 0 – 0.25 s post tap-event). **B. Averaged delta peak topographies** (top: closest to behavioural tapping rate, bottom: high delta). **C. Per-participant correlations between the tapping-evoked topography and delta-peak topography.** Left: Pearson’s r values, right: p-values, FDR corrected across participants. The horizontal dashed line indicates the cut-off of p = 0.05.

### 2.12. Analyses of oscillatory bursts

We performed a confirmatory analysis (depicted in **Figure 9**) to further assess whether the spectral peaks we identified reflect endogenous periodicities in the time-domain data, which are likely not fully stationary. To this end, we quantified **oscillatory bursts** using the cycle-by-cycle toolbox [76]. To our knowledge, this method has so far not been used for frequencies as low as delta. To test whether the sensors identified as having a spectral peak in a given canonical frequency band really show stronger periodic activity, we assessed the time-domain signal of those sensors for bursts. The data were band-pass filtered between 0.5 and 4.5 Hz prior to performing the analysis. The filter band was chosen somewhat larger than the band of interest to allow for transition bands. The cycle-by-cycle algorithm labels peaks and troughs in the filtered time-domain signals, and then computes several statistics to identify truly periodic episodes, defining oscillatory bursts. The threshold parameters used to detect episodes with bursts were adapted for the delta band: amplitude fraction threshold = 0.05, amplitude consistency threshold = 0.4, period consistency threshold = 0.4, monotonicity threshold = 0.95, minimum number of cycles = 2.

**Figure 9.**
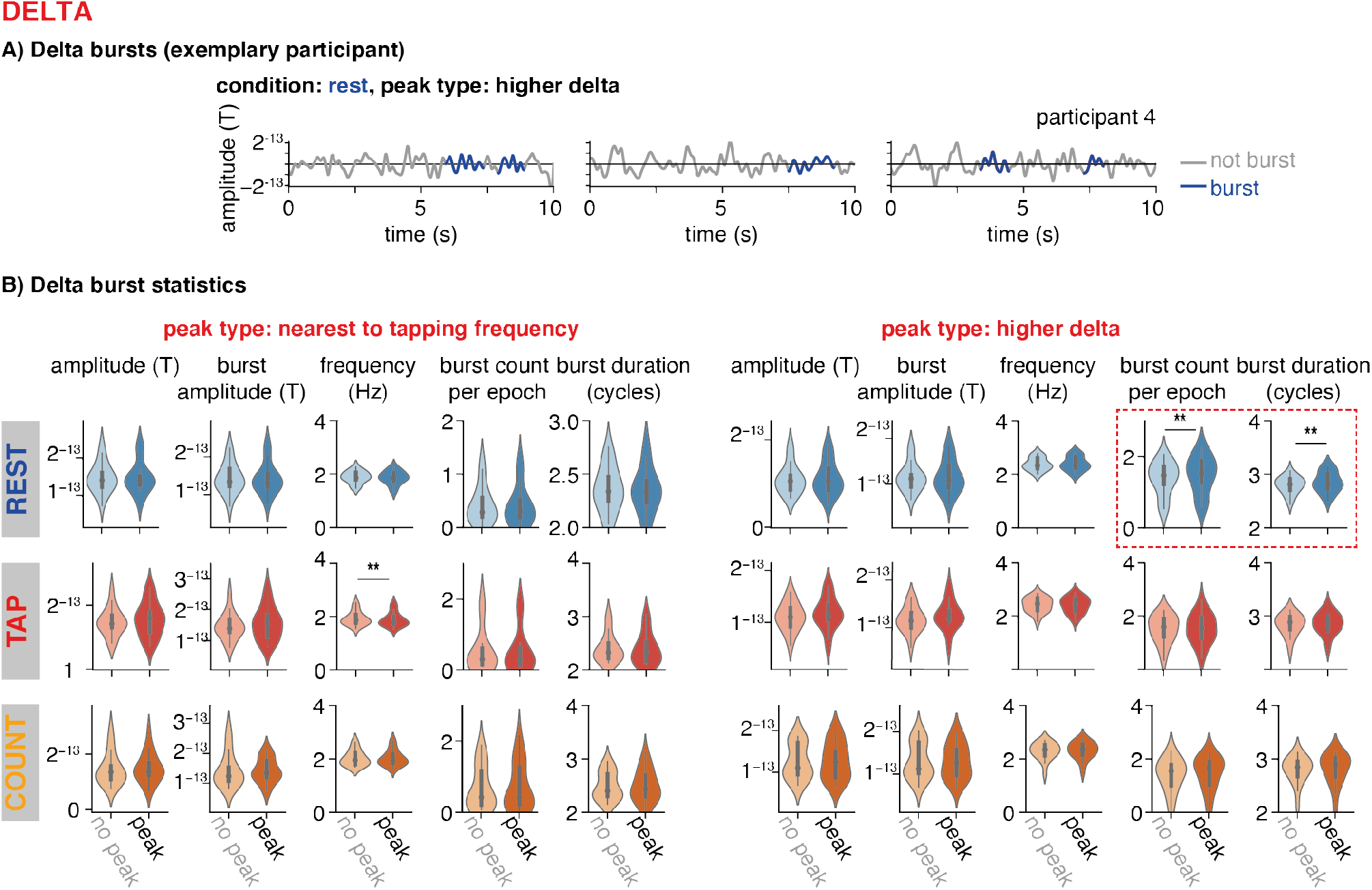
**Analysis of oscillatory bursts, delta band**, comparing the statistics resulting from the cycle-by-cycle analysis (amplitude, burst amplitude, frequency, burst count, burst duration) at the sensors that were identified to have a peak in the delta frequency band (dark violins) versus a randomly sampled selection of other sensors (light violins). The columns show the two peak types identified in the delta range: tapping rate and higher delta. The rows show the three conditions, rest, tap, count. Significantly more and longer bursts were found in the peak-sensors for the higher delta band in the resting condition, as indicated by the red dotted square.

To confirm whether the peaks detected in the spectral analyses described above reflect endogenous periodicities, we computed the by-cycle analysis on the signals from the sensors identified as belonging to a given cluster (sensors with peak), and on a random selection of sensors of the same number not belonging to the cluster (no peak, averaged over 50 repetitions of random selection for stability). We ran this analysis for all three conditions, and per participant (**Figure 9,** see **Figure S6** for the other canonical frequency bands; thresholds for the theta/ alpha/ beta bands: amplitude fraction threshold = 0.3, amplitude consistency threshold = 0.6, period consistency threshold = 0.6, monotonicity threshold = 0.95). The detected burst episodes were summarized by the average burst amplitude, burst period, duration, and number of bursts, on which we performed paired samples t-tests between the two types of sensors (with peak and without peak, threshold p < 0.01).

## 3. Results

### 3.1. Behavioural results

#### 3.1.1 Tapping and counting rates

Individual tapping rates were estimated through spectral analyses of the button press times as illustrated in **Figure 4A** (mean rate = 1.17 Hz, SD = 0.6 Hz, **Figure 4B** shows the distributions). Individual covert counting rates were estimated by dividing the final count number reported by the participant by the duration of the task (mean = 0.93 Hz, SD = 0.3 Hz, measure available for 17 out of 18 participants, **Figure 4B**). Overt tapping and covert counting rates were found in the same range (no significant difference: T(16) = -1.21, p = 0.45), and did not correlate within participants (Pearson’s r(15) = -0.28, p = 0.28, **Figure 4C**).

#### 3.1.2. Subjective duration estimates

The relative subjective duration estimates (estimated duration in seconds divided by the run’s duration) were 1.14 s (SD = 0.40) and 0.91 s (SD = 0.34) for the tapping and counting runs, respectively. Relative duration estimates did not correlate significantly within participants between the tapping and counting runs (Pearson’s r(16) = 0.33, p = 0.19, not shown). In the tapping run, there was no significant correlation between the overt individual tapping rate and the relative duration estimate (r(16) = −0.20, p = 0.42), in the counting run, there was a strong correlation between the covert counting rate and the estimated duration (r(15) = 0.94, p < 0.01). We also tested whether the spectral peak frequencies (reported in detail below) correlated with the subjective duration estimates for the respective runs, in all frequency bands, but no significant correlations were found (p > 0.09).

### 3.2. Spectral peak detection

Spectral peaks were identified for all participants in all conditions and frequency bands, including the delta range (see **Table 1** for peak frequencies, power, and the number of sensors per peak cluster). As shown in **Figure 5**, the group topography, computed by averaging across participants’ topographies at their individual delta peaks, shows a typical profile of frontocentral activity, likely emerging from motor and auditory regions. Importantly, and in particular for the delta band, the topographies of oscillatory activity averaged across the canonical frequency band resembled the topographies of the 1/f activity (see **Figure 2**), while the averaged topographies at individually detected peak frequencies show distinct frontocentral profiles (**Figures 3, 5**). Individual peak sensors (depicted by coloured markers on the topographies, Fig. 5) show that the sensors at which the strongest delta power was detected varies across participants, but generally cluster in accordance with the average topographies.

As described in Section 2.7, three types of peaks were selected from the clustering procedure: one being **closest to the individual’s behavioural tapping rate**, separately identified per condition (average peak frequencies in resting, tapping, counting runs: 1.25 Hz (SD = 0.43), 1.37 Hz (0.5), 1.56 Hz (0.56), respectively; average power in rest, tap, count: 3 dB (SD = 0.64), 3.69 dB (SD = 1), 2.75 dB (SD = 0.58), respectively, **see Figure 6, Table 1**). The second peak was selected as being ***above* the individual’s tapping rate** (average peak frequencies in resting, tapping, counting runs: 2.29 Hz (SD = 0.5), 2.35 Hz (SD = 0.53), 2.09 (SD = 0.57), respectively; average power in rest, tap, count: 3.15 dB (SD = 1.04), 3.31 dB (SD = 1.06), 3.06 dB (SD = 1.29)). The third peak was selected as **closest to the individual’s *counting* rate** (average peak frequencies in resting, tapping, counting runs: 1.22 Hz (SD = 0.32), 1.08 Hz (SD = 0.32), 1.15 (SD = 0.33), respectively; average power in rest, tap, count: 3.22 dB (SD = 0.73), 3.33 dB (SD = 0.96), 2.97 dB (SD = 0.65)).

No significant differences in delta peak frequency (**Figure 6A, Table 2**), were found between the three conditions (two comparisons marginal: p = 0.023, 0.056, all other p-values > 0.21), at all three peaks. The peak frequencies selected close to the *counting rate* were not significantly different from the peaks selected close to the *tapping rate*, although the difference was marginally significant for the data recorded from the counting condition (rest: T(16) = -0.04, p = 0.97; tap: T(16) = -1.64, p = 0.12; count: T(16) = -2.41, p = 0.03).

Peak power at the individual tapping rate was significantly higher in the tapping condition (paired samples t-test, tapping > resting: T(17) = 3.0, p < 0.01; tapping > counting: T(17) = 3.49, p < 0.01, **Figure 6B**). We did not find a similar pattern for peaks identified by being close to the individual counting rate, that is no enhanced peak power in the counting condition (all p > 0.4). The number of sensors showing a peak at the detected frequency was marginally larger in the tapping condition compared to resting (tapping > resting: T(17) = 2.52, p = 0.02, **Figure 6C**).

### 3.3. Delta peaks: correlation analyses

As intended by the peak selection procedure, the spectral peaks identified as closest to the behavioural tapping rate showed strong correlations with the tapping rate in all conditions (**Figure 7A**). While this correlation is trivially related to our peak selection, it does reflect that in every participant a spectral peak could be found in the vicinity of the spontaneous tapping rate. The spectral peaks identified as closest to the behavioural counting rate showed no significant correlation with the counting rate in the resting and counting condition, but in the tapping condition (**Figure 7B**), suggesting that the spectral peaks in the counting runs did not result from covert counting, but rather overlapped with the tapping peaks.

We then calculated the residual frequency differences after accounting for the correlation between *tapping rates* and peak frequencies (by subtracting the frequency values predicted by the linear fit). The residuals were significantly smaller in the tapping condition compared to rest and count (tap < rest: T(17) = 3.38, p < 0.01; tap < count: T(17) = 6.28, p < 0.01, **Figure 7C**), indicating that stronger activity around the individual behavioural tapping rate was present in the MEG recordings of the tapping condition. We performed the same analysis for the peaks selected as closest to the behavioural counting frequency, but the residuals frequency difference between the spectral peaks and counting rate were also smallest in tapping and not counting (**Figure 7D**), confirming once more that the selected peaks did not distinctively result from covert counting.

Only for the peaks identified by tapping behavior, we then further tested the residuals for correlations across conditions, hypothesizing that an endogenous oscillation present in all conditions would result in relatively stable peak frequencies (and hence stable residuals from the behavioural tapping rate) across conditions. However, no significant correlations were found (**Figure 7E,** tapping and counting: r(16) = 0.44, p = 0.07; tapping and resting: r(16) = - 0.18, p = 0.48; counting and resting: r(16) = 0.05, p = 0.84).

In an additional control analysis (not shown), we also extracted the first peak, defined as the cluster with the largest number of sensors from the delta band instead of dividing peaks by their closeness to the behavioural tapping or counting rate. No significant correlations were found between the strongest spectral peak in the tapping condition and tapping behaviour (r(16) = 0.09, p = 0.72) or the strongest peak in the counting condition and counting behaviour (r(15) = -0.43, p = 0.12). The absence of a significant correlation, especially in the case of tapping, can be explained by the existence of multiple heterogeneous peaks per participant and condition. The clustering approach found a peak close to the behavioural tapping rate for each participant, but this was not necessarily the strongest cluster.

### 3.4. Comparison of tapping evoked activity with spectral peaks

To explore whether the spectral peaks observed in the MEG activity during tapping reflect the periodic occurrence of the tapping evoked response, or an ongoing oscillation, we compared the topographies of the two neural signatures (**Figure 8**). For the spectral peak at the tapping rate, correlations between the topographies of the tapping evoked response (**Figure 8A**) and the peak topography were found in the majority of participants (13/ 18, **Figure 8B/C top**). The topography of the higher delta peak correlated with the topography of the tapping evoked response in eleven participants (**Figure 8B/C bottom**). While it was expected to see a correlation between the peak at the tapping rate and the tapping evoked response, it is more surprising that the topographies of the higher delta peaks also correlate with the tapping evoked response. Two explanations can be thought of: either the higher peak also partially reflects the periodically occurring evoked response, or it reflects an oscillation in overlapping brain regions.

### 3.5. Oscillatory bursts

In a confirmatory analysis, we compared different statistics of oscillatory burst episodes in the delta band (burst amplitude, frequency, count, and duration) between sensors at which a spectral peak had been identified and a random selection of sensors without a peak (see **Figure 9**). For the delta peaks close to the individual tapping rate, the only significant difference was found in burst frequency in the tapping condition with a higher frequency at the no-peak sensors (T(17) = 3.32, p < 0.01). In the resting condition, we found significantly higher burst counts (T(17) = 3.5, p < 0.01) and longer burst duration (more cycles, T(17) = 3.29, p < 0.01, **Figure 9B**) for the higher delta band, in the sensors with peaks (marginally more bursts counted also in the counting condition: T(17) = 2.06, p = 0.06). The finding that the overall amplitude did not differ at the sensors with peaks compared to no peaks can be explained by the broader frequency range used in this analysis, while as reported above, the peak detection procedure showed variability in frequency across participants.

### 3.6. Theta-, alpha-, beta-bands

For validation of the approach, we also ran this analyses on the other canonical frequency bands (see **supplementary Figure 5**), and found strong differences in the alpha band for all burst statistics in the resting condition (all p < 0.01), marginally more and longer bursts in the low beta band (p = 0.05, 0.03), and marginally longer burst episodes in the high beta band (p = 0.05), for the tapping condition only.

## 4. Discussion

### 4.1. Summary

Here, we assessed whether endogenous delta oscillations can be observed in non-invasively recorded human brain dynamics. As per the current state of the art, we define oscillations by a narrow peak in the power spectral density [71]. We analysed an existing MEG dataset [75], from which we selected three runs during which participants either rested, or engaged in rhythmic behaviour through spontaneous finger tapping (overt engagement of the motor system) and silent counting (covert engagement of the auditory-motor system), and concurrent prospective timing. Dedicated signal processing techniques were applied, involving a multi-step procedure for removal of confounding artifacts in the delta frequency range (cardiac activity), and separation of aperiodic (1/f) components of the power spectrum from periodic components. Our results clearly show that narrow spectral peaks can be observed in the delta band in human MEG but require refined signal processing techniques. The peaks were heterogenous across conditions and participants, and partially overlapped with evoked activity, but intrinsic periodicities were identified during rest.

### 4.2. Locally oscillating neural populations

A first important observation was that several spectral peaks were detected in the delta band per participant and condition (**Figures 3, 5**). Peaks were heterogenous with respect to the peak frequency, topographical distribution across sensors, and strength, i.e., the number of sensors showing a peak at a given frequency. Crucially, on average, or in a group statistical approach, no coherent peaks could be found, while individual power spectra do show signatures of oscillatory activity. This suggests that what we pick up in the M/EEG is the synchronous signal from relatively small, and spatially local neuronal populations. This point has long been made by Hari et al., who reported "…that synchronous activity of 1% of cells in a cortical area of 1 cm^2^ would determine 96.5% of the total signal" [89], caption of Fig.12, p.60. The current literature often considers neural oscillations as a rather widespread phenomenon, but recent studies have started to separate local oscillators. For example, one study investigated local alpha oscillations in auditory areas in humans with depth electrodes implanted in temporal regions and reported two distinct oscillators in primary and secondary auditory areas [90], see also [91]. Another study used high density EEG to show that the seemingly paradoxical observation of theta oscillations surfacing during both rest and cognitive effort reflects separable neural dynamics [92]. The sparseness and variability of oscillating populations and their differential contributions to the recorded signals likely exacerbate divergences between existing human M/EEG studies, and should thus be taken into account more thoroughly. We hope that the work presented here provides an example and some possible guidelines in this direction, especially for the case of slow oscillations.

### 4.3. Do spectral peaks reflect endogenous periodicities?

A critical question is whether the spectral peaks that we identified truly qualify as endogenous oscillations, versus reflect periodically occurring evoked activity. The peaks found in the tapping condition were closer to the individual tapping rate, and stronger in power compared to the resting and counting conditions (**Figure 6, 7**), reflecting enhanced delta activity when participants engaged in tapping. This is in line with previous studies that linked self-paced tapping to neural dynamics in the delta frequency range [93], [94], but did not distinguish between evoked responses and oscillatory dynamics. Here, we observed the strongest delta activity at central and frontal sensors, but the peak topographies correlated with the topographies of the tapping evoked response in the majority of participants (**Figure 8**). This suggests that the spectral peak in the vicinity of the individual tapping rate reflects at least in part the periodically occurring sensorimotor evoked responses [59], [87], and neural signatures of spontaneous movement [95]. We thus cannot claim that the peaks in the tapping condition reflect endogenously periodic activity. This questions the commonly used practice to search for spectral peaks when analysing neural oscillations, at least in the presence of other potential sources of periodic activity. Future functional studies will be necessary to compare delta activity in MEG recordings across different tasks.

To further investigate whether the spectral peaks reflect intrinsic periodicities, we performed a confirmatory analysis to classify non-stationary oscillatory delta band bursts in the time domain (**Figure 9**) [76]. We found significantly enhanced oscillatory activity in the high delta band during resting, namely more and longer burst episodes in the sensors that forming the peak cluster. While the application of this method to low frequency oscillation such as delta should be further validated, our finding provides preliminary evidence for spontaneous delta band oscillations in non-invasive human MEG recordings during rest.

The topography of the high delta peaks during rest (**Figure 5**, top right panel), confirmed as oscillatory in the additional analysis, suggests motor, frontal, and possibly also auditory generators, in line with previous studies [9], [39], [96]. Notably, these oscillatory bursts occurred at frequencies above the behavioural tapping rate, pointing towards independency between tapping-evoked delta activity and endogenous oscillations. It might be the case that an activation of the auditory system by sensory inputs would have engaged internal oscillators in auditory regions more strongly, as endogenous delta oscillations have previously been reported in primary auditory cortices [8], an assumption that can be tested in future work.

We also examined the hypothesis that spontaneous rhythmic activity is orchestrated by a stable internal frequency, i.e. a propensity of the relevant neural populations to oscillate at this frequency, by correlating peaks in tapping and counting behaviour and delta peaks across conditions [66]–[69], [97]. The behavioural frequencies measured in tapping (1.17 Hz) and counting (0.91 Hz) were clearly in the delta range, albeit somewhat lower than the frequencies of the spectral peaks in the MEG data (avg. tapping peak: 1.39 Hz, avg. peak above tapping: 2.43 Hz, avg. counting peak. 1.24 Hz), and lower than the preferred frequency recently reported for auditory-motor synchronization 1.7 Hz, [69]. There was no correlation of peak frequencies across conditions after controlling for the peak selection frequencies (**Figure 7C**), which would have been expected if the peaks observed in the three conditions resulted from a stable neural oscillation. The behavioural frequencies measured in tapping and counting also did not correlate within individuals (**Figure 4**). Thus, the assumption that spontaneous rhythmic motor behaviour (overt as in tapping, or covert as in counting) reflects the frequency of a common neural oscillator could not be confirmed here. However, it is possible that the small N underlying the correlation analyses prevented such an observation [85].

### 4.4. Comparison between the present results and previous invasive studies

As mentioned before, the existence of endogenous delta oscillations and their roles for cognitive processing are derived from seminal animal work. In humans, the availability of invasive recordings is very limited, but several studies provided important insights from intracranial recordings in epilepsy patients. Our results align with these studies in several aspects, notably the observation of delta activities in various brain areas, and the variability of the oscillatory patterns over time. In a task-based study, Besle et al. [31] show indirect evidence for endogenous delta oscillations by observing phase resets in a wide-spread network including motor cortex, orbitofrontal cortex, angular gyrus, and parietal regions (70% of recording sites), suggesting endogenous delta oscillations occur in many brain regions. In the same line, a seminal investigation by Halgren et al. [9] addressed the generators of endogenous delta activity across cortical areas (frontal, parietal, temporal) and cortical layers. Delta (and theta) activity was prominently observed during sleep and wakefulness (rest) at all recording sites, with local generators in superficial cortical layers. Our results align with this work and underline the necessity to go beyond Fourier-based methods, to assess to what extent peaks observed in low frequency bands reflect intrinsic periodicities. In an analysis restricted to auditory areas, Neymotin and al. [98] identified ’oscillatory events’ occurring regularly in the resting state data. Both the local field potentials (non-human primates) and intracranial EEG recordings (humans) during rest showed prominent delta activity. Yet, the oscillations were not stationary, and occur for 3 cycles on average in the delta band, which nicely matches with our time-domain analyses.

In sum, there is clear evidence for endogenous delta oscillations in various areas of the human brain, mostly supported by local invasive recordings, and in line with the host of task-based studies describing activities in the delta band. As evidence converges that delta oscillations are local and not stationary, it will be crucial to take into account the biological properties of these dynamics in the study of cognitive dynamics in humans.

### 4.5. Theta, alpha, and beta bands

For validation of the analysis pipeline, and comparison, the same analyses were run on three other canonical frequency bands (see **supplementary Figures**): theta (4 – 7 Hz), alpha (8 – 12 Hz), and beta (15 – 30 Hz). Most participants had several peaks per frequency band with considerable variation in the peak frequencies. Further examination of the time domain signal for intrinsic periodicities using the cycle-by-cycle method showed that besides the delta band, the only clearly oscillatory activity was found in the alpha band, confirming previous studies [75], [99].

While recently new methods have been proposed for the analysis of neural oscillations [76], [98], [100], most have been validated in the alpha band but see [101] for a refined examination of neural dynamics in the the beta band. In the alpha range, dominant spectral peaks can be easily identified, and periodic activity can be spotted in the time domain with the naked eye. It is currently an open question whether those methods perform less well when detecting oscillations at slower frequencies, or whether dynamics in other frequency bands differ so strongly in their characteristics from the prominent alpha activity that it is even questionable whether they qualify as endogenous oscillations. A better understanding of the characteristics of neural oscillations beyond stationary sinusoids is required to move forward on these questions [102].

Surprisingly, we observed no enhanced beta power in the tapping condition, previously reported during auditory-motor synchronization [103]–[105]. Beta oscillations have also been reported to reflect explicit duration estimates [106]–[108], and implicit non-rhythmic temporal predictions [30], [32], [109]. As recently reported, beta oscillations occur as short bursts at specific moments in time, rather than as stationary oscillations [110], [111], which might explain why we did not observe increased power during the whole tapping run. This further supports the notion that Fourier based methods do not account well for the non-stationarities present in the brain dynamics and thus miss the presence of oscillations. In line with this assumption, the additional time domain analyses indicated at least marginally more and longer burst counts in beta during tapping (supplementary **Figure S6**, bottom left panel).

### 4.6. Outlook

Delta oscillations seem indicative of a critical balance between widely synchronized and local activity, where too much synchronization is a signature of pathological brain activity [112], but local oscillations appear crucial for engaging with a particular task. The presented results argue for the necessity characterize each delta oscillatory profiles per individual, namely in terms of peak frequency and topography. These individual profiles, obtained based on the methodological guidelines we outlined, can be used as the basis for future studies interested in the functional relevance of slow neural oscillations. The outlined procedures can be used to derive a spatio-spectral filter (or localizer), serving to extract the signal of interest, for instance when interested in mechanisms of synchronization to external rhythms [113], or memory consolidation, functionally linked to delta oscillations. This approach can also be applied in the assessment of clinical populations that show characteristic alterations in delta oscillations [9], [114], [115], for instance in schizophrenia were increased delta oscillatory power is observed over frontal regions. The approach proposed here will hopefully help to promote a more unified reporting of the physiological characteristics of delta oscillations relevant to a particular task, or pathology, to eventually enable meta-analytic approaches as currently available for alpha oscillations [116].

### 4.7. Limitations

Here, we present a first attempt to assess endogenous delta oscillations in non-invasive MEG recordings in humans. While our work shows that spectral peaks can be observed in the delta band, we would not want to assume that those peaks reflect endogenous oscillations in all experimental conditions. One particular limitation was the rather short recordings we used (2 min per condition and participant). Given the observation that oscillatory episodes might be transient [98], [101] short recordings might have resulted in a low signal to noise ratio. Longer datasets should be screened for endogenous delta oscillations with the same methods to observe their consistency over time, and their relationship with mental states and behavioural tasks in awake human participants. Longer datasets would also allow to address phase-amplitude coupling between delta oscillations and higher band power, as previously observed [8], [9]. Furthermore, the small sample size (N = 18) might have limited our chances to find significant differences and correlations [85], which should be considered in interpreting the results.

### 4.8. Conclusions

Here we examined whether endogenous delta oscillations can be observed in non-invasive MEG recordings in humans, by analysing human resting state recordings. To test whether spontaneous rhythmic behaviours incite an otherwise silent internal oscillator, we also selected conditions in which participants engaged in spontaneous finger tapping and silent counting. Narrow spectral peaks in the delta frequency range were found in all conditions, but additional analyses targeting non-stationary oscillations in the time domain showed that only the resting state data warranted an interpretation of endogenously periodic neural dynamics. We hope that the novel set of analyses steps and results presented here will foster a more detailed investigation of spontaneous oscillations in low frequency bands in humans, and thus contribute to better standards in characterizing the physiological signals underlying a hypothesized functional mechanism, and enable comparison across studies in healthy and clinical population.

### 4.9. Data and Code Availability

Raw magnetoencephalography recordings, as well as the pre-processed epochs for all three runs per participant will be made available on the Open Science Framework together with the complete analysis pipeline (python code), and the behavioural data. The currently private repository will be made public upon acceptance of the manuscript for publication.

## 7. Additional information

### 7.1 Acknowledgements

We would like to thank the UNIACT staff at NeuroSpin for their help with participant recruiting and the MEG recordings.

### 7.2. Author contributions

Original study design: LA, VvW

Data collection: LA

Conceptualization of present study: SH

Data curation and analyses: HG, SH

Writing first draft: HG, SH

Editing: HG, LA, VvW, SH

Resources: CEA, VvW

Funding for the project: VvW

All authors reviewed and approved the manuscript.

