## Supplementary Materials for "Characterizing endogenous delta oscillations in human MEG"

### 1. Supplementary Materials

#### 1.1. Spectral peak detection: theta, alpha, beta bands

Individual spectral peaks detected in the theta, alpha, and beta bands from an exemplary participant are shown in **Figures S1, S2, S3**. The average peak topographies are depicted in **Figure S4** (theta, alpha, beta bands). The theta peaks occurred most strongly at frontal sensors (**Figure S4**, green), alpha at parietal (**Figure S4**, purple), and beta at central sensors (**Figure S4**, orange). Low beta power was marginally higher in the tapping compared to the counting condition (T(17) = 2.34, p = 0.03; **Figure S5**), but not when compared to resting (T(17) = 0.97, p = 0.34). No other condition differences were observed.

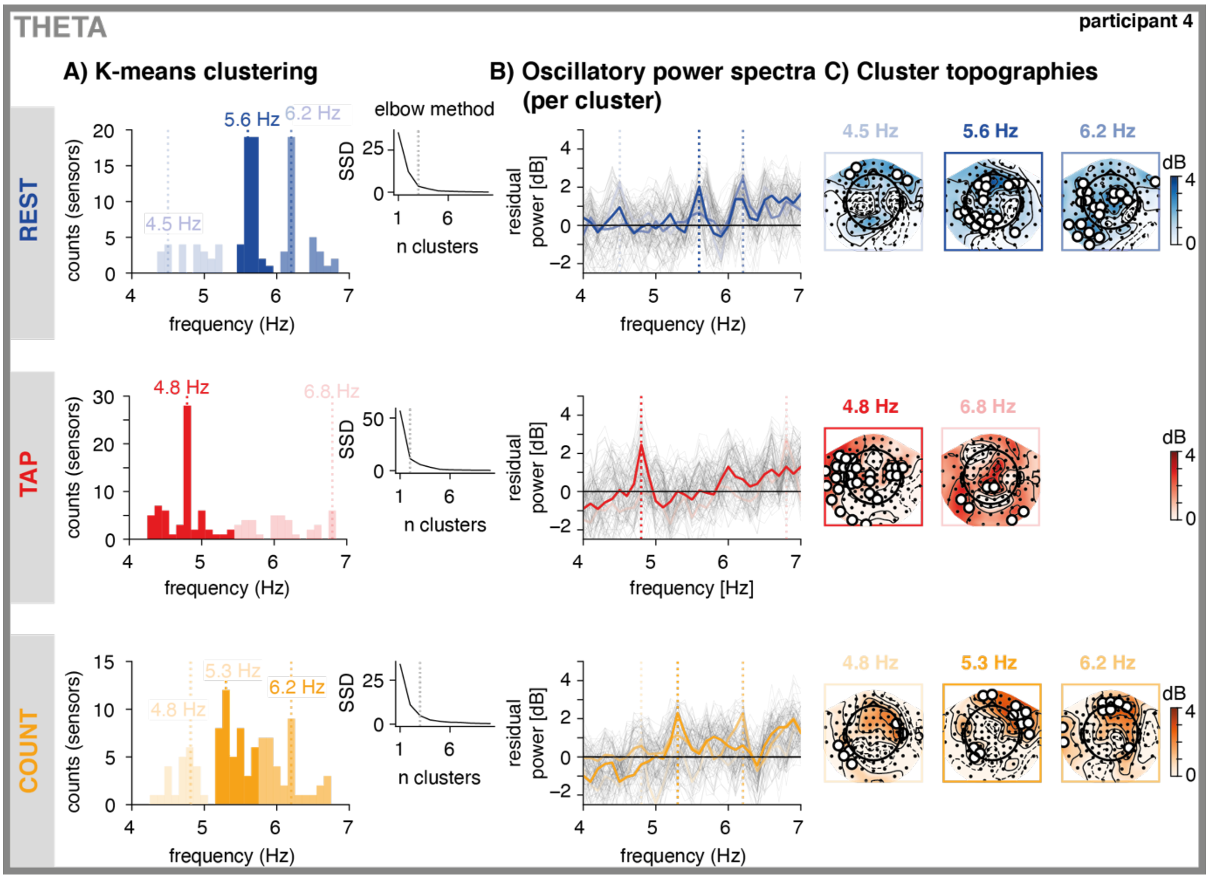


**Figure S1. Theta band, peak selection in an exemplary participant. A. K-means clustering.** Per condition (rest, tap, count, depicted as rows and blue, red, and orange), spectral peaks identified per sensor were clustered using k-means (histograms; the brightness codes for cluster order (brighter = higher), with the strongest clusters in saturated colours). The vertical dashed lines indicate the peak frequency per cluster (defined as the mode). The number of clusters were determined from the data by identifying an 'elbow' in the goodness of fit (SSD, the sum of square distances from each point to its assigned cluster centre, see inset). **B: Oscillatory power spectra**, extracted from all sensors (grey lines), and the sensors in each cluster (blue, red, and orange lines). The vertical dashed lines depict the peak frequency of each cluster. **C: Cluster topographies**. Power distribution across all sensors at the cluster's peak frequency (± 0.1 Hz). The white circles depict sensors selected as part of the cluster.

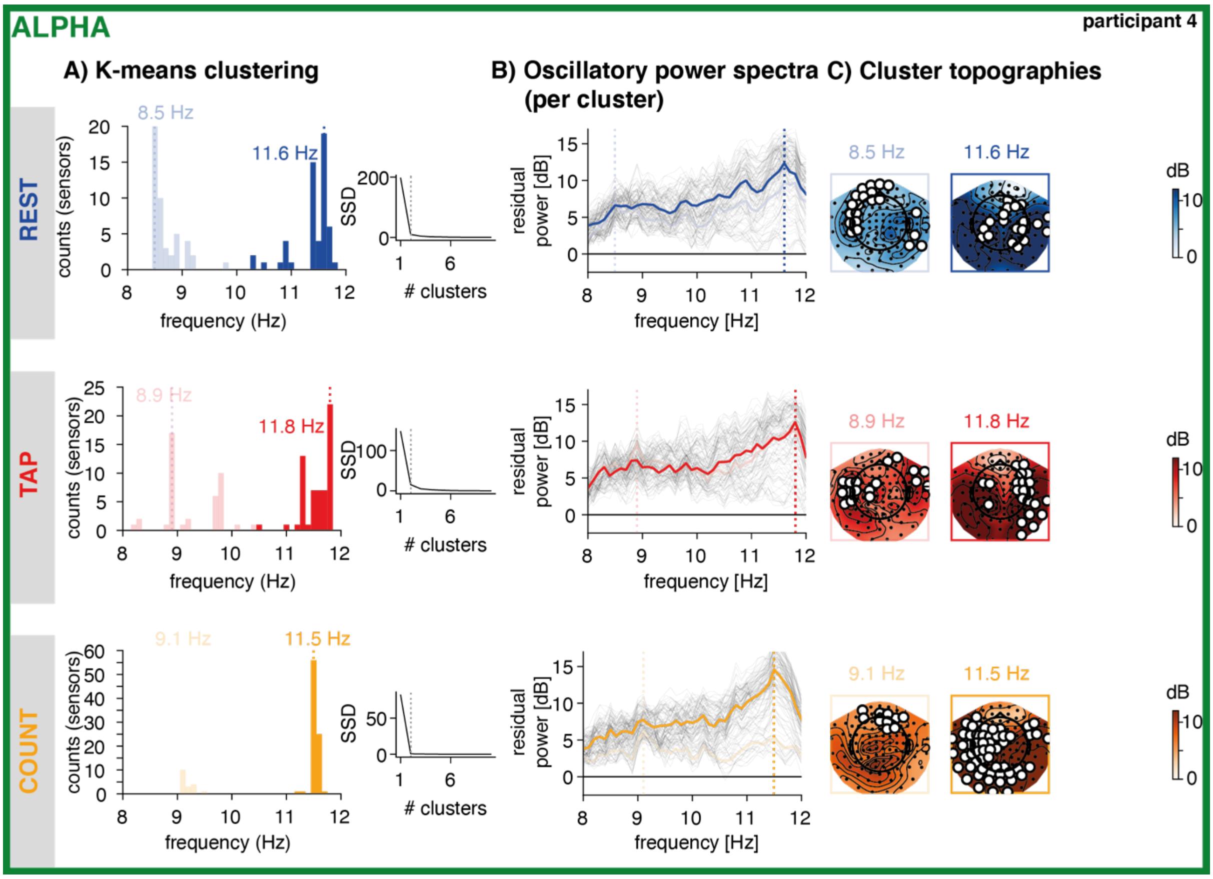


**Figure S2. Alpha band, peak selection in an exemplary participant**. **A. K-means clustering.** Per condition (rest, tap, count, depicted as rows and blue, red, and orange), spectral peaks identified per sensor were clustered using k-means (histograms; the brightness codes for cluster order (brighter = higher), with the strongest clusters in saturated colours). The vertical dashed lines indicate the peak frequency per cluster (defined as the mode). The number of clusters were determined from the data by identifying an 'elbow' in the goodness of fit (SSD, the sum of square distances from each point to its assigned cluster centre, see inset). **B: Oscillatory power spectra**, extracted from all sensors (grey lines), and the sensors in each cluster (blue, red, and orange lines). The vertical dashed lines depict the peak frequency of each cluster. **C: Cluster topographies**. Power distribution across all sensors at the cluster's peak frequency (± 0.1 Hz). The white circles depict sensors selected as part of the cluster.

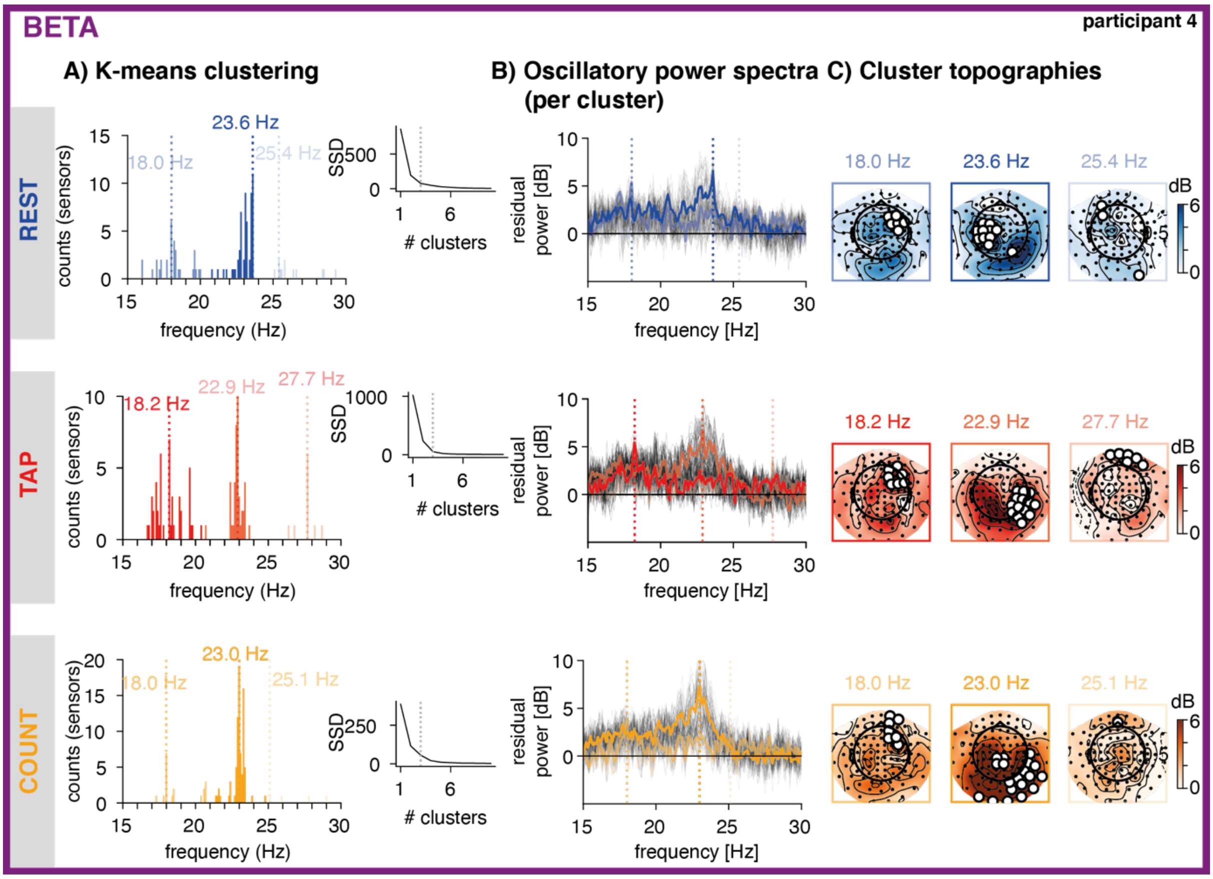


**Figure S3. Beta band, peak selection in an exemplary participant. A. K-means clustering.** Per condition (rest, tap, count, depicted as rows and blue, red, and orange), spectral peaks identified per sensor were clustered using k-means (histograms; the brightness codes for cluster order (brighter = higher), with the strongest clusters in saturated colours). The vertical dashed lines indicate the peak frequency per cluster (defined as the mode). The number of clusters were determined from the data by identifying an 'elbow' in the goodness of fit (SSD, the sum of square distances from each point to its assigned cluster centre, see inset). **B: Oscillatory power spectra**, extracted from all sensors (grey lines), and the sensors in each cluster (blue, red, and orange lines). The vertical dashed lines depict the peak frequency of each cluster. **C: Cluster topographies**. Power distribution across all sensors at the cluster's peak frequency (± 0.1 Hz). The white circles depict sensors selected as part of the cluster.

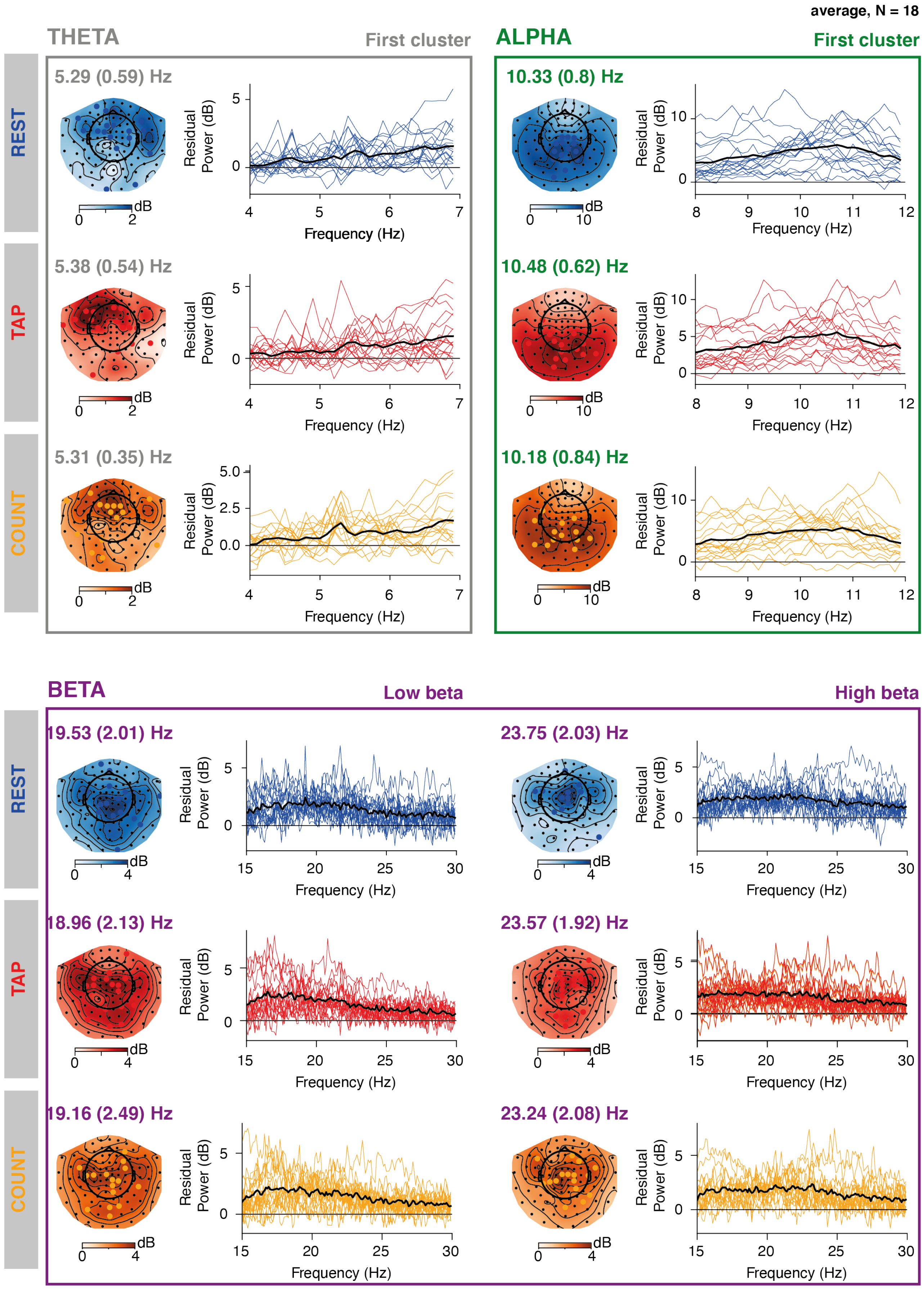


**Figure S4. Result of the peak sorting procedure, theta, alpha, beta bands.** In the theta (grey square, with conditions in blue (rest), red (tap), and orange (count) and alpha (green square, top right) bands, only the first cluster (defined by the largest number of sensors having a peak at that frequency) was extracted, in the beta band, two clusters were extracted divided into low (purple square, bottom left) and high beta band activity (purple square, bottom right). Peak sorting was done per participant and condition (tap, rest, count, depicted in the rows). The topographies depict the power distribution at the individually identified peak frequencies, averaged across participants. The values above the squares indicate the averaged peak frequency and standard deviation across participants Individual oscillatory power spectral density at the cluster's sensors (coloured lines) and average across participants (black line).

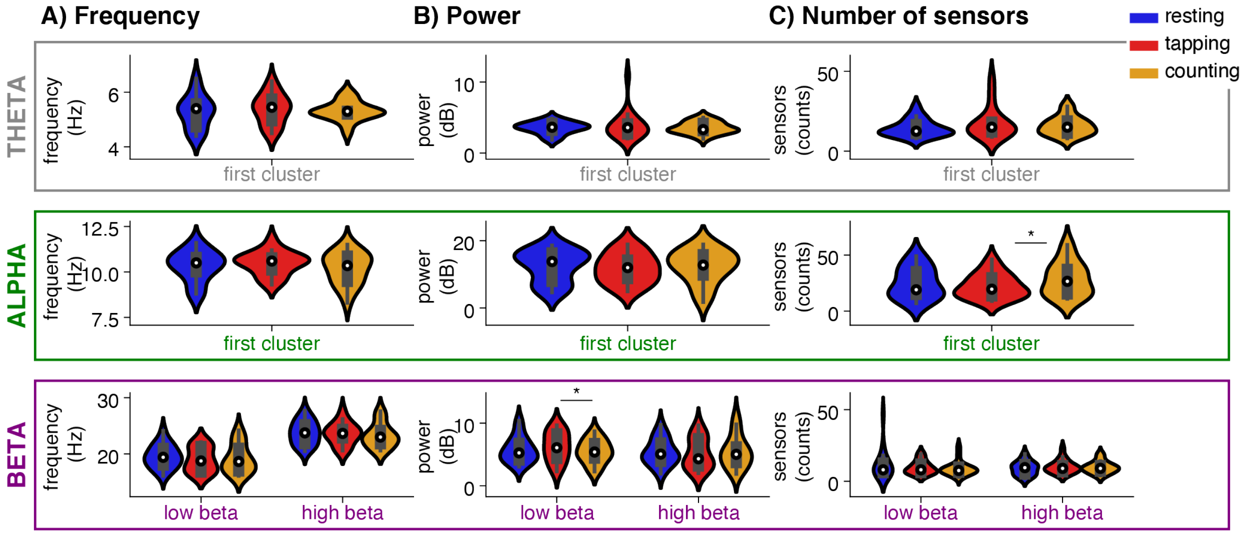


**Figure S5. Peak frequency and power per canonical band and condition. A: Peak frequency.** Violin plots depict the distribution (kernel density estimation) of peak frequencies across participants. White dots: median. Thick grey bar: interquartile range. **Theta, alpha:** peak frequency distributions of the first clusters. **Beta:** peak frequency distribution of the low and high beta activity. **B. Peak power.** As in A. **C. Number of sensors in cluster.** As in A. Significance values are indicated as follows: * p < =0.05; ** p < 0.01.
